# Microbial Colonisation of Polyethylene in Offshore Marine Environments: Insights from the Southern and South Atlantic Oceans

**DOI:** 10.1101/2025.05.18.654703

**Authors:** Priscilla Carrillo-Barragán, Gabriel Erni Cassola, Patricia Burkhardt-Holm

**Affiliations:** The Man-Society-Environment Programme, Department of Environmental Sciences, University of Basel, Switzerland; Department of Plant and Microbial Biology (IPMB), University of Zurich, Zurich, Switzerland

**Keywords:** Plastisphere, plastic marine debris, microbial colonisation, plastic biodegradation, plastic biofilms, South Atlantic Ocean

## Abstract

Plastic debris is a pervasive environmental pollutant with polyethylene (PE) among the most abundant floating polymers in marine environments. While microbial colonisation of marine plastics has been extensively documented, studies predominantly focus on Northern Hemisphere coastal waters, leaving microbial dynamics in remote Southern Hemisphere oceanic regions poorly characterised. In this study, we employed ribosomal amplicon sequencing and shotgun metagenomics to characterise microbial colonisation patterns on PE films deployed during two oceanographic transects across the Southern Ocean and South Atlantic Ocean over 14 days of incubation. We evaluated environmental factors including geographic location, incubation regime (indoor/outdoor), and exposure conditions (UV radiation, temperature, and salinity) as potential drivers shaping microbial communities. Our results demonstrate clear differentiation in microbial community structure driven primarily by transect location and environmental conditions, rather than material type. Dominant taxa identified included Pseudomonadota known for hydrocarbon degrading capabilities and Cyanobacteria associated with phototrophic traits. Metagenomic analyses revealed enrichment of functional pathways linked to biofilm formation, hydrocarbon degradation, and iron metabolism. This study expands the current understanding of microbial colonisation processes on marine plastics, highlighting environmental factors influencing early plastisphere community structure and functional potential in understudied oceanic regions.

## 1. Introduction

Global plastic pollution is widely recognised as a major environmental threat to marine ecosystems due to its widespread distribution and persistence (Barnes et al., 2009; Thompson et al., 2009). On sea surfaces, buoyant polymer types such as polyethylene (PE), the most produced and discarded plastic polymer worldwide (Geyer et al., 2017), are found at higher concentrations (Eriksen et al., 2014). Floating PE and other plastic marine debris (PMD) become rapidly colonised by marine microorganisms, forming biofilms known as the “plastisphere” (Zettler et al., 2013). Such microbial colonisation affects the fate of these anthropogenic materials by potentially influencing their buoyancy, degradation rates, biota uptake, and capacity to transport putative pathogens and associated contaminants across marine ecosystems (Amaral-Zettler et al., 2020; Carrillo-Barragan et al., 2023; Kooi and Koelmans, 2019).

Biofouling by larger calcifying organisms, such as barnacles and bryozoans, has been identified as a critical driver of sinking plastic debris (Fazey and Ryan, 2016). However, the fate of smaller plastic debris that cannot support macroscopic biofouling remains poorly understood. Specifically, the role of microbial communities, often the sole colonisers of small-sized PMD, on the dynamics of these smaller plastics is understudied. Recent work highlights that early colonising microbial communities on plastic surfaces significantly influence subsequent community development and functionality, emphasising the importance of characterising early biofilm formation stages (Bos et al., 2023).

Marine microbial communities colonising plastics are known to differ from the surrounding seawater, shaped by environmental factors such as temperature, salinity, ultraviolet radiation, and geographical region (Kesy et al., 2019; Ogonowski et al., 2018; Schlundt et al., 2020; Wright et al., 2020; Zeghal et al., 2024). Yet, current knowledge primarily originates from studies in coastal and northern hemisphere waters, leaving significant gaps regarding plastisphere communities in remote offshore environments, particularly in the Southern Hemisphere (Amaral-Zettler et al., 2015; Oberbeckmann et al., 2018; Scales et al., 2021; Schlundt et al., 2020).

The Southern Ocean and South Atlantic Ocean represent unique ecological environments with distinct environmental pressures, such as colder temperatures, heightened UV exposure, and specific nutrient conditions (Convey and Peck, 2019; Pörtner, 2006). Antarctic and sub-Antarctic ecosystems, characterised by simple food webs include species adapted to extreme isolation, pronounced seasonality, and limited energy availability (Convey and Peck, 2019; Pörtner, 2006). Such organisms display physiological traits that heighten their vulnerability to pollutants, including slow metabolism, limited detoxification capabilities, and reliance on lipid reserves (Leistenschneider et al., 2022; Leuenberger et al., 2024). Therefore, understanding microbial colonisation on PMD in these regions is particularly important, as microbes influence pollutant dynamics, plastic buoyancy, potential degradation pathways and PMD biota uptake, with direct implications for regional ecological health (Carrillo-Barragan et al., 2023; Convey and Peck, 2019; Wright et al., 2020; Zadjelovic et al., 2022).

Despite emerging evidence that plastic pollution has reached remote Antarctic waters, impacting local organisms across multiple trophic levels (Leuenberger et al., 2024; Wilkie Johnston et al., 2023), microbial colonisation processes of PMD in these regions remain scarcely studied. Recognising this critical gap, we hypothesise that distinct environmental pressures in Southern Hemisphere offshore waters shape early microbial colonisation patterns on PE plastics, driving both taxonomic differentiation and functional succession in plastisphere communities.

In this study, we investigate microbial community dynamics on PE films deployed during two oceanic transects in the Southern Ocean (SO) and South Atlantic Ocean (SAO). Specifically, we aim to determine: (a) whether early stage plastisphere community structures differ significantly between these geographically distinct oceanic regions, and (b) how key ecosystem roles (generalist, specialist, and transient taxa) develop during initial microbial colonisation, focusing on metabolic functions relevant to motility, adhesion, hydrocarbon degradation, and nitrogen fixation. To address these objectives, we employed 16S rRNA amplicon sequencing and shotgun metagenomics, offering detailed insights into microbial succession and functional potentials during the early colonisation of polyethylene in these underexplored oceanic environments.

## 2. Methods

The Polarstern experiment was conducted to simulate the early prokaryotic colonisation of drifting PE films in surface marine waters along offshore transects of the Atlantic Ocean at different time points. Incubated plastic samples and glass slides controls were deployed and harvested during two consecutive cruise research expeditions aboard the RV Polarstern icebreaker 1) expedition PS129 from Cape Town to Punta Arenas (03.03.2022 – 28.04.2022), 2) expedition PS130/1 from Punta Arenas to Las Palmas (30.04.2022 – 22.05.2022) (Hoppema, 2023). Our sampling locations along these transects are herewith referred as Transect 1 (Tr1) and Transect 2 (Tr2), respectively (Figure 1). The distance between the last sampling point during Tr1 and the first at Tr2 was of approximately 2,2224.5 Km, while the distances between sampling stations within transects were in the range of 630.2 Km to 1078.8 Km and 2559.0 Km to 3319.9 Km, for Tr1 and Tr2, respectively.

**Figure 1.**
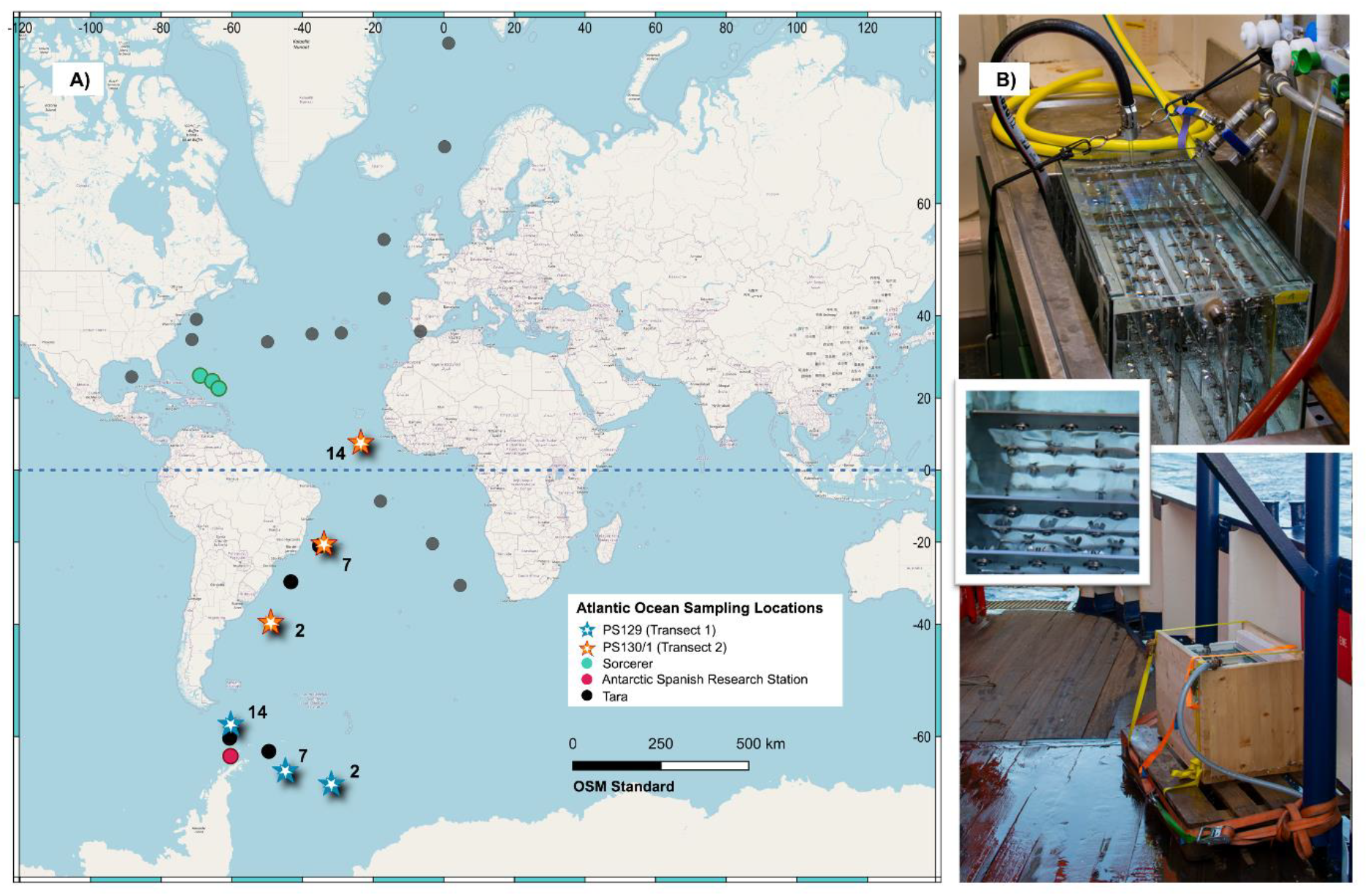
A) World map with i) sampling points for expeditions PS129 (blue stars) and PS130/1 (orange stars) in this study; ii) sampling points of the Sorcerer II (green circles) and iii) sampling station for the Antarctic Spanish Research station (pink circle), expedition and incubations which are relevant to this work, respectively; iv) sampling points for microbial analysis of the Tara Oceans expedition (2010 - 2011) in the Atlantic Ocean (black circles). The blue dotted line marks the Equator. B) Incubation set up with PE films and glass slides suspended in stainless steel frames inside tanks located inside an aquarium and outside on the ship deck.

### 2.1 Sample preparation and incubations set up

Samples consisted of i) non-weathered and ii) UV pre-weathered (254 nm, 15 W/m^2^ for 14 days) PE film cut outs (25 × 75 mm) from plastic carrier bags, and microscopy glass slides which were used as controls and treated as PE films. All plastic films and glass slides were pre-cleaned in 10% bleach then rinsed in filter-sterilised distilled water for 30 min, followed by three rinses with laboratory grade deionized water (previously filter-sterilised). Both sample types were attached to stainless steel frames in replicates of 4 and placed in pre-cleaned (10% bleach then rinsed in filter-sterilised distilled water) glass fish tanks (w x h x d= 30 × 36 × 60 cm) filled with seawater continuously pumped from the ship’s Klaus Union Sealex Centrifugal Pump (stainless steel piping) for flow through incubations in all incubation types (Figure 1).

Individual fish tanks with samples submerged in seawater (60 L) were placed in parallel on a) the ship deck and exposed to natural sunlight, and b) inside one of the ship’s aquaria, with nearly all collections taking place in the morning between 9 – 12 hrs.

Samples (x4 replicates) were incubated underway and harvested incrementally across the cruise track. Two independent transects (Tr1 and Tr2) were selected to compare the early stages of PE films prokaryotic colonisation (Figure 1). Due to extremely stormy weather, only ship’s aquaria incubations were conducted for Tr1. As reference, the Tara Ocean expedition sampling locations for planktonic prokaryotic samples in the Atlantic Ocean are included in the map on Figure 1, highlighting the novelty of the samples used in the present study.

### 2.2 Sample collection and storage

After a pre-determined number of days (2, 7, and 14 days), samples were removed with sterile forceps from the stainless-steel rack and rinsed with filter sterilised seawater. Samples were then immediately placed in Falcon tubes containing 40 mL of filtered sterilised seawater and sonicated for 5 minutes at maximum frequency (6000 rcf). The sonication process was repeated three times per sample. Plastic films and glass slides were further rinsed with filter sterilised seawater and then removed from the Falcon tubes. Falcon tubes were centrifuged for 20 minutes at 8000 rcf (centrifuge SIGMA 2K15 Zent 1). The supernatant was discarded, and the cell pellet resuspended in 350 µL of lysing solution (solution MBL, DNeasy PowerBiofilm Kit). The suspension was transferred to 1.5 mL centrifuge tube and stored at - 20°C until further analysis.

### 2.3 DNA extraction, library preparation, and 16S RNA sequencing

Pellets from PE films and glass slides replicates for day 2, 7 and 14 of each incubation set up were submitted to the NU-OMICS DNA Sequencing Research Facility (Northumbria University, Newcastle, UK). Genomic DNA was isolated from samples using the DNeasy PowerBiofilm Kit (Qiagen, UK) following the manufacturer’s protocol. A procedural negative control (nuclease free water) and a mock community standard as positive control (D6305 ZymoBIOMICS, UK) were processed with each batch of samples. The Earth Microbiome Project (EMP) (Thompson et al., 2017) 16S Illumina Amplicon Protocol was followed, including the use of the 16S SSU rRNA V4 region primer set (515F–806R) for PCR amplification. The pooled library was prepared and validated using a Quant-iT PicoGreen dsDNA Assay Kit (Invitrogen, UK) and a 2100 Bioanalyzer (Agilent Genomics, UK). Illumina MiSeq sequencing was done following a V2 2 × 250 bp chemistry (Illumina, UK).

### 2.4 16S rRNA sequence library processing and community composition analyses

Raw sequencing data was processed into amplicon sequence variants (ASVs) using the DADA2 pipeline (Callahan et al., 2016) within RStudio (R version 4.4.1) (Posit team, 2025). Quality filtering, trimming (240 bp for forward reads and 160 for reverse reads), and error detection (EE = 2) was followed by sequence dereplication, alignment, merging of for-ward and reverse reads, and chimera removal, resulting in 2981 ASVs detected across 111 samples. Taxonomy was assigned using the SILVA 138 reference database (Quast et al., 2012). The ASV table and metadata were further analysed using the Phyloseq R package (version 1.48.0) (McMurdie and Holmes, 2014).

### 2.5 Statistical Analyses

Non-parametric statistical tests were used to evaluate ASV data. Richness (ASVs counts), diversity (Inverse Simpson’s index), and evenness (Simpson’s Evenness) were calculated to assess α-diversity. Microbial community composition was analysed using a Bray-Curtis dissimilarity matrix derived from square root-transformed ASV abundance data. A Non-metric Multidimensional Scaling (NMDS) ordination plot was used to visualise clustering among the microbial communities growing on the different incubation treatments. Differences between groups were assessed using Permutation Multivariate Analysis of Variance (PERMANOVA, adonis2 function, 999 permutations) to quantify the effects of time (day of sampling), geographic region (Transect), and material (PE, wPE, glass) on community structure, and Analysis of Similarity (ANOSIM) to validate group separation based on rank similarities. Similarity Percentage Analysis (SIMPER) was employed to identify taxa contributing most to differences between groups, focusing on the top 10 contributors per comparison. Unclassified taxa were aggregated to the lowest available taxonomic level for interpretability. All analyses were performed in R using the vegan (Oksanen, 2008) and phyloseq (McMurdie and Holmes, 2014) packages.

### 2.6 Profiling of shotgun metagenomic sequences

To infer the metabolic potential of prokaryotic communities growing in the different incubation treatments, samples of interest were selected for further shotgun metagenomic sequencing (n = 75). Samples were processed at the NU-OMICS Sequencing Facility (Northumbria University, UK) using the Illumina NextSeq 500 (New England Biolabs).

Raw sequences were analysed for taxonomic and functional profiling using Kraken2, Bracken, and HUMAnN3 as previously described (Franzosa et al., 2018). For each sample, paired-ended reads were quality-trimmed using Trim Galore to remove adapter sequences and low-quality bases using (Franzosa et al., 2018). Taxonomic classification of trimmed reads was performed using Kraken2 followed by Bracken v2.9 for species-level abundance estimation against the core_nt database (inclusive of GenBank, RefSeq, TPA and PDB) from the AWS Public Dataset Programme (AWS, 2024). Bracken was executed with a read length of 100, a confidence threshold of 10, and classification set at the species level. The HUMAnN3 pipeline (Franzosa et al., 2018) was used for profiling the presence/absence and abundance of the pathways and gene families in each incubation treatment. The output files from HUMAnN3 were summarised in R to compute mean abundances and the standard deviation within each incubation group. Gene families of interest were matched against the UniProt ID mapping tool to infer their functions.

Raw amplicon and metagenomic sequencing data are openly accessible at the NCBI SRA archive (BioProject ID: PRJNA1195827), and biosamples accession numbers as well as other sequencing processing metadata are available at the Zenodo Data repository DOI 10.5281/zenodo.15437519.

### 2.8 Comparison with the literature

A reference table (S1, Bos et al., 2023) compiling studies on microbial communities growing in plastic in marine environments from 2013 to 2021 was expanded to include relevant research from 2017 to October 2024 (Supplementary Table 2 “ST2”). Original peer-reviewed publications were compiled in August 2024 and revised in October 2024 searching in the Scopus database the keywords “plastic” or “plastic marine debris” or “microplastic” and “aquatic” or “marine” and “biofilm” or “plastisphere” or “epiplastic”. From the 719 documents found for the selected period, 469 focused on marine plastics of which only 25 investigated the Atlantic Ocean’s plastic biofilm communities.

ST2 includes the sequencing method and sample collection location for each study. Notably, from the total 51 studies in ST2 (Bos et al compilation plus the references added by the present study), only 6 of these (including the present work) analysed the plastisphere community composition of South Atlantic, Southern Ocean, or Antarctic samples. At the moment of writing, no previous study had conducted a shotgun metagenomic analysis (WGS) of such communities.

Due to their methodological or location similarities, suitable samples from Bos and colleagues during the expedition R/V Sorcerer II (Bos et al., 2023) and from Monràs-Riera (Monràs-Riera et al., 2024) Antarctic microcosms’ incubations, are directly compared with our results (ST1). Briefly, the expedition R/V Sorcerer II samples of interest (days 3, 5 and 7, ST1) were collected during the transect from West Palm Beach, Florida to Marigot on the island of St. Martin (19–26 March 2017). These 7 day-long shipboard incubations consisted of incubating high-density polyethylene (HDPE) jug punches and pre-industrial PE pellets (Bos et al., 2023). Additionally, Monràs-Riera and collaborators (Monràs-Riera et al., 2024) examined the plastisphere of UV-weathered PE pellets in continuously flowing Antarctic seawater at the Antarctic Spanish Research Station (ASRS) in Livingston Island (South Shetlands, Antarctica). The team subsampled over a month long (January–March 2022) outdoor incubations, including at 2, 5, and 12 days - samples of interest in the present study- and evaluated bacterial community physical distribution (Scanning Electron Microscopy), and composition (16S rRNA amplicon sequencing and qPCR).

## 3. Results & Discussion

### 3.1 Profile of early-colonising prokaryotic communities extracted from different materials in relation to environmental factors

An initial analysis of the prokaryotic communities across different incubation treatments over time was conducted based on ASVs, focusing on alpha diversity indices (Fig. 2A) and richness in relation to temperature and salinity (Fig. 2B), environmental factors deemed as key influences on microbial community structure (Oberbeckmann et al., 2018; Sunagawa et al., 2020).

**Figure 2.**
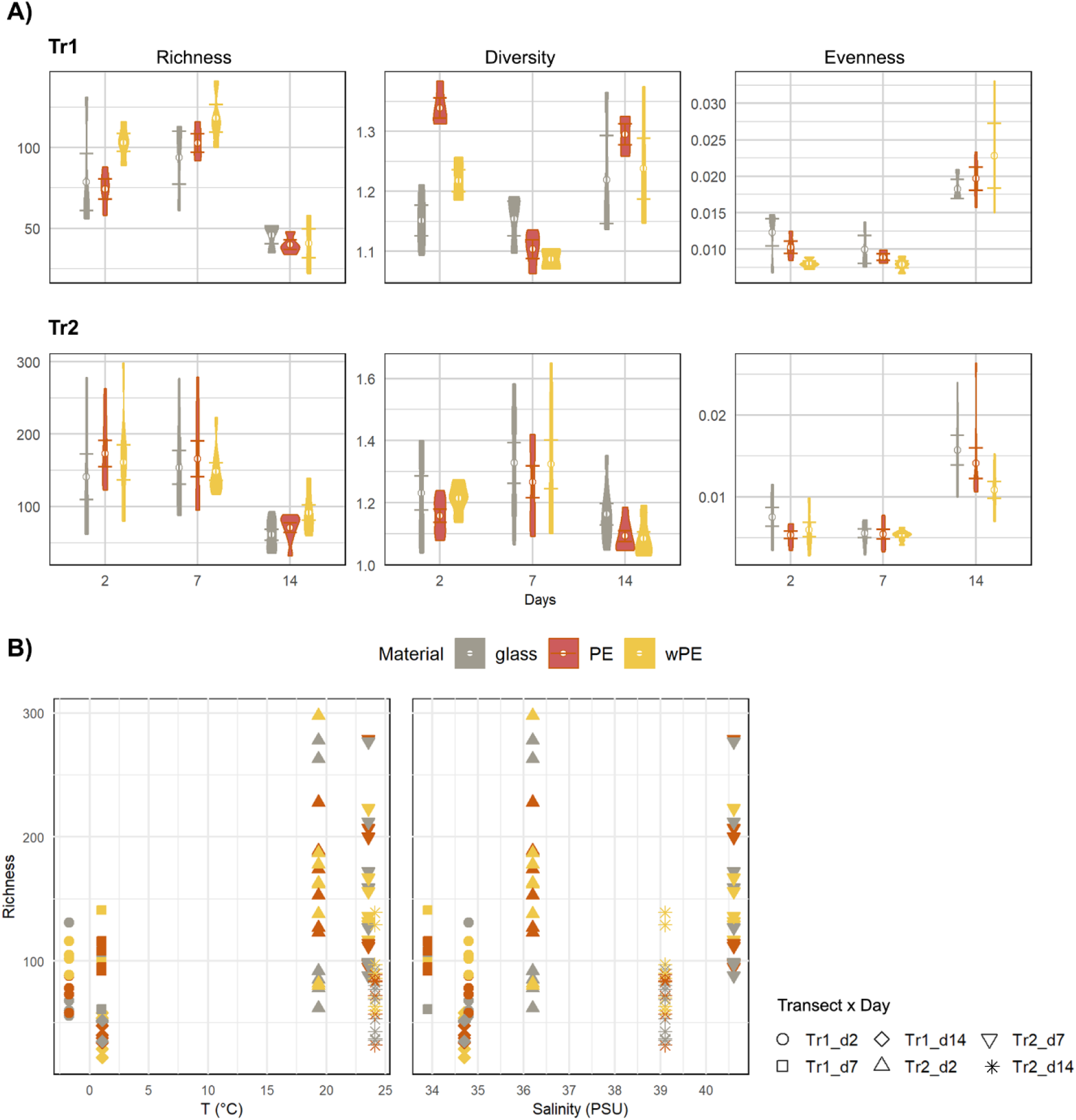
A) Alpha diversity indices of communities colonising the different materials tested over time. B) Richness as a function of water temperature (°C) and Salinity (PSU) at the different sampling stations. Tr1 and Tr2 = Transect 1 and Transect 2. d2, d7, d14 = sampling day 2, day 7, day 14. Error bars represent standard error of the mean.

Our results agree with previous observations of low alpha diversity in pre-weathered PE (Erni-Cassola et al., 2020). Nevertheless, ASVs richness was significantly higher in Tr2 incubations than in Tr1 (2-way ANOVA, p < 4.8 e-07), though diversity did not differ significantly between the two transect (2-way ANOVA, p = 0.8606). The latter is highlighted by the broad variations in diversity values observed within replicates (Fig. 2A). Collection day significantly impacted richness, with a sharp decline by day 14 (Tukey’s test, p < 1e-07), suggesting temporal shifts potentially linked to environmental factors or community maturation (Erni-Cassola et al., 2020; Zeghal et al., 2024). Evenness increased progressively over time (p < 2e-16), with Tr2 incubations exhibiting consistently higher richness but lower evenness than Tr1 (Tukey’s test, p < 2e-09 for evenness between transects), likely due to differences in species dominance or in the process of community maturation. Material type (PE, wPE, glass) did not significantly affect richness, diversity, or evenness within transects. These results emphasise the prominent roles of geographic region and time variability in structuring community richness and evenness and agree with previous studies where material type had a minimal effect on microbial community assembly on plastic substrates (Monràs-Riera et al., 2024; Zeghal et al., 2024).

We further tested the relationship between latitude and richness (Fig. 2B), to evaluate if plastic colonisation would follow the generally agreed observations of richness increasing with ocean water temperature (4 – 12 °C) in intermediate latitudes (Sunagawa et al., 2015). Similarly, salinity has been regarded as a primary environmental factor influencing microbial community structure (Guan et al., 2023; Telesh et al., 2013). Thus, we also evaluated if the relatively narrow range of salinity changes across latitudes experienced in our incubations (34.8 to 40.6 PSU, 3.4% to 4.0%) would have an impact in the richness of these plastisphere communities.

Despite the relatively wide range in temperatures (< -1 °C – 24 °C) between Sothern Ocean and South Atlantic Ocean waters, the richness displayed by PE communities did not show a significant relationship with temperature (Kendall’s Tau = 0.0119, p = 0.8672). In contrast, salinity had a mildly significant positive relationship on richness (Kendall’s Tau = 0.2423, p = 0.0007). These results suggest that the incubated PE colonising communities display different biogeographic patterns to those reported in previous studies evaluating planktonic prokaryotic communities (Guan et al., 2023; Sunagawa et al., 2020; Telesh et al., 2013). These differences highlight how attachment to a surface can influence the structure of microbial communities, as seen in other studies of the plastisphere in marine environments (Oberbeckmann et al., 2018; Schlundt et al., 2020). Variations within each group (Fig. 2) may have also resulted from small-scale environmental differences or random colonisation events, both of which are known to shape plastic-associated microbial communities (Kesy et al., 2019).

#### 3.1.2 Differential community dynamics on PE, wPE, and glass early colonising prokaryotic communities across geographical regions

Microbial β-diversity was estimated using a Bray-Curtis dissimilarity matrix and visualised with a non-metric multidimensional scaling (nMDS) plot (Figure 3). The nMDS (stress value 0.1) showed clear clustering based on transect and day of sampling, also displaying distinct ordination between aquarium and on deck incubations. ANOSIM analysis further examined microbial community structure across these different factors, revealing significant community separation by day (R = 0.063, P = 0.001), transect (R = 0.811, P = 0.001), and incubation location (inside the aquarium versus outside on the boat deck; R = 0.294, P = 0.001). However, ANOSIM indicated non-significant differences in community composition based on material type (R = 0.021, P = 0.103), suggesting minimal rank-based separation by material. In contrast, PERMANOVA (P(perm) = 0.001) showed that material significantly influenced microbial community structure (Pseudo-F = 4.464), along with day of sample collection per transect (Pseudo-F = 17.300), incubation location (Pseudo-F = 31.979), and transect (Pseudo-F = 52.195). Together, these analyses suggested that although material type did not create strong rank-based separation, it nonetheless accounted for a significant, albeit mild, proportion of overall variance in microbial community composition.

**Figure 3.**
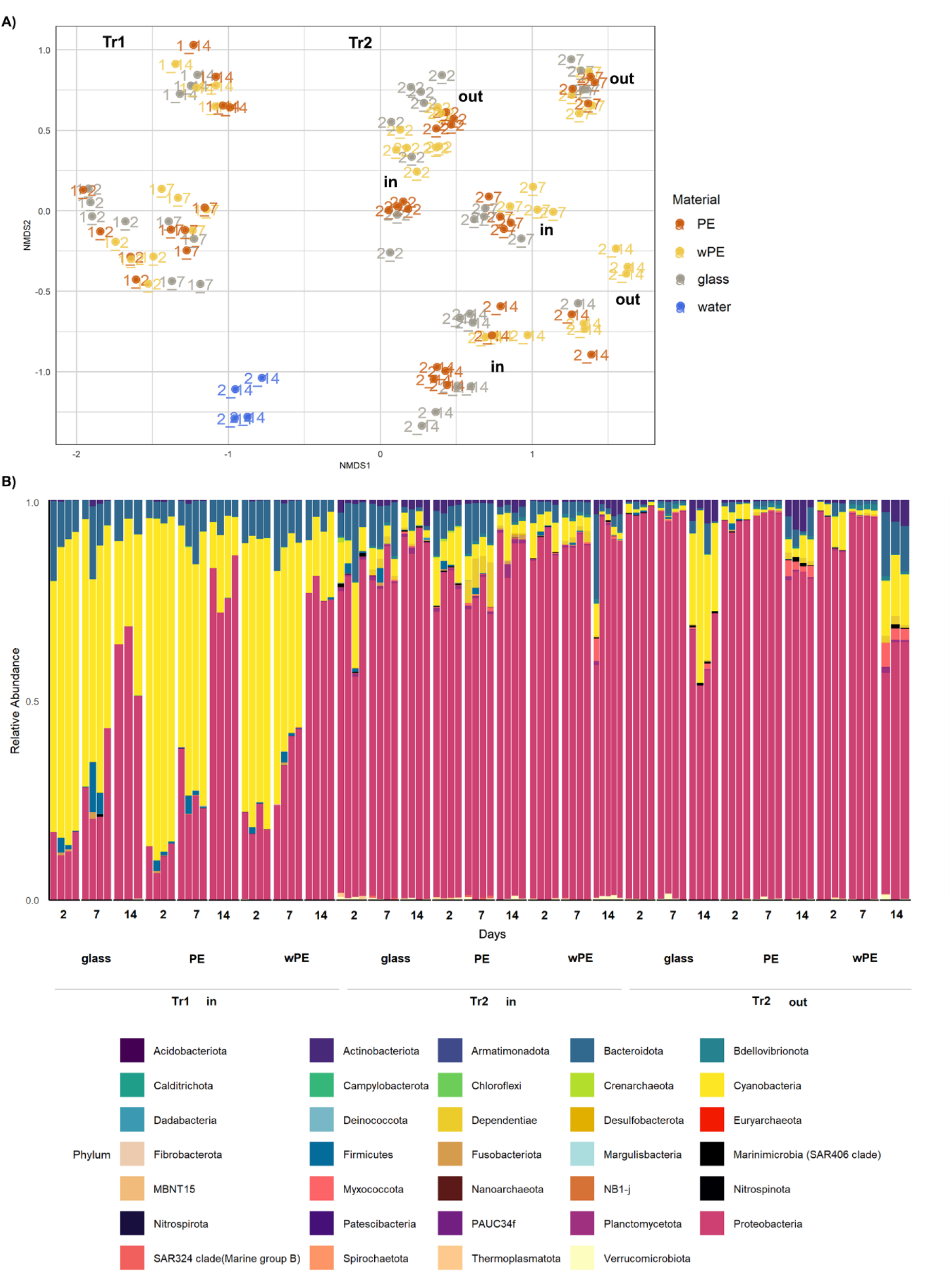
A) nMDS plot of onboard incubation communities at different sampling times (days 2, 7, 14) over two independent transects of the Southern and South Atlantic Oceans. Each circle represents a replicate of a material in given transect and day (denoted “1_2” for Transect 1 day 2). Place of onboard incubation is denoted by “in” for aquarium incubations and “out” for ship deck incubations. B) Relative abundance of prokaryotic Phyla growing in the different incubation treatments. Transects are denoted as Tr1 and Tr2 on plots A) and B).

The relative abundance of ASVs across treatment groups (Fig.3) revealed the presence of 34 phyla, and a general dominance of Cyanobacteria, Pseudomonadota, and Bacteroidota during the experiment (Fig. 2B). These results confirmed that the low alpha diversity displayed by the communities in all incubation treatments were due to a small number of abundant species that drove the differentiation of the communities. Notably, distinct temporal variation was observed between transects: Tr1 incubations, initially dominated by Cyanobacteria (82.7% ± 1.9%), shifted towards Pseudomonadota (79.1% ± 6.3%) enrichment by the end of the incubation period. In contrast, Tr2 exhibited consistent dominance by Pseudomonadota throughout the experiment (ranging between 61.3 % ± 5.2 % - 96.7 % ± 1.1 %), with a mild increase in Cyanobacteria in the glass (from 1.2 % ± 0.8 % to 28.5 % ± 9.9 %) and wPE (from 5.5 % ± 3.5 % to 13.9 ± 1.3 %) treatments incubated outdoors over time. These findings align with previous studies, confirming Cyanobacteria, Pseudomonadota, and Bacteroidota as recurring colonisers of marine plastics (Aguila-Torres et al., 2022; Bryant et al., 2016; Roager and Sonnenschein, 2019). Furthermore, the dominance of Cyanobacteria in Tr1 is supported by datalogger measurements from the Polarstern expedition in the Southern Ocean (Hoppmann et al., 2023), where phycocyanin levels peaked at 444 cells/ m^3^, indicating localised cyanobacterial blooms.

For further characterisation, a pair-wise analysis of dissimilarity (SIMPER) between plastisphere communities showed that communities incubated on all materials experienced temporal shifts in dominant taxa, with notable differences between incubation inside an aquarium versus outside on the ship deck (Figure 4). For instance, the increased relative abundances of of taxa such as Rhodobacteraceae, *Cycloclasticus*, and *Alteromonas*, specially under outside incubation conditions, further highlight the synergistic effect between photooxidative degradation and biodegradation, as previously noted by Albertsson et al. (1987). This is also demonstrated by the selective enrichment, in Tr2 outside incubations, of Parvibaculales, an order of the alpha proteobacteria known to colonise plastic surfaces (Debroas et al., 2017; Roager and Sonnenschein, 2019), including genera able to use synthetic surfactants as carbon source (Schleheck et al., 2011).

**Figure 4.**
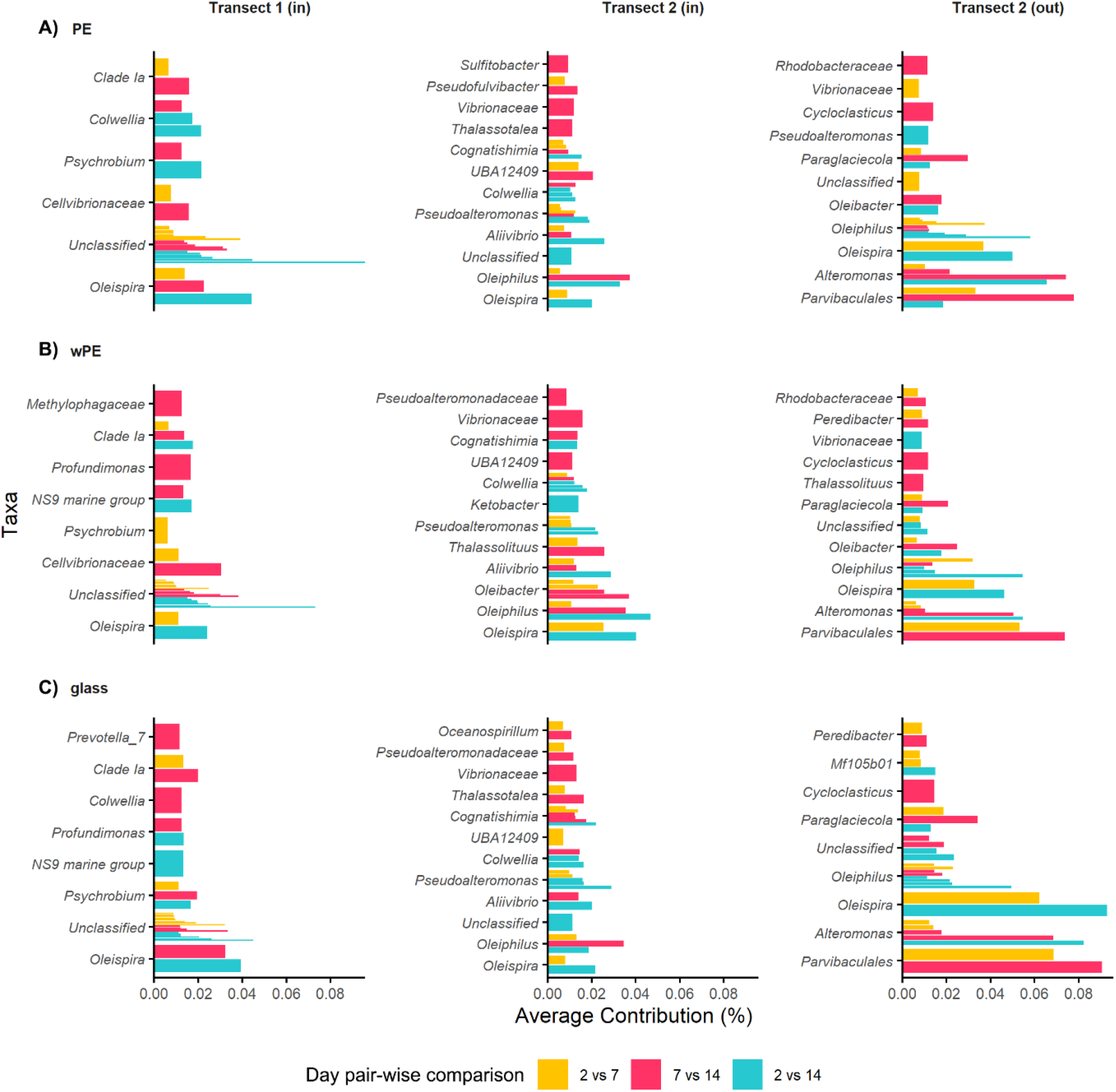
SIMPER day pair-wise comparison of ASV top taxa contributing to the dissimilarity between communities in each material over time. (in), (out): inside aquarium or outside on ship deck incubation, respectively. Unclassified: prokaryotic sequences unclassified in the Silva database v1.38.

Among the top unique contributors to dissimilarities between transect communities, the transitionary enrichment of the gamma proteobacteria *Methylophagaceae* in wPE while the plastisphere community was still maturing (days 7-14) in Tr1, suggests the labile one-carbon compounds this common to polar surface waters halophilic methylotrophic (Ramachandran et al., 2021) were easily available during this time period and readily consumed by the end of the incubation period.

However, the materials were mostly colonised by similar taxa. These results align with those of Bos et al. (2023), Monras-Riera et al. (2024), and others (Erni-Cassola et al., 2020; Harrison et al., 2014) finding that early microbial colonisers on PE (Bos et al., 2023) and wPE (Monràs-Riera et al., 2024), include genera like *Oleispira, Oleiphilus, Colwellia*, and *Pseudoalteromonas* (ST1, Fig. S1), all known for hydrocarbon degradation and biofilm formation. The present study also observed these genera as top drivers of community differences for both, PE and wPE, being highly abundant at early stages of colonisation and decreasing by the end of the incubation period (Fig.4, days 2 to 14) across all incubation conditions. Our findings demonstrate that polyethylene surfaces, like other abiotic substrates such as glass, can create niches for microbial specialist functions, including hydrocarbonoclastic prokaryotes, across a broad range of geographical regions, including the South Atlantic and Southern Ocean waters.

### 3.2 Species-level taxonomy profiling of polyethylene growing communities in South and Southern Atlantic waters

The initial Bracken processing of shotgun metagenomic sequencing data showed high variability in the success of taxonomic classification and filtering thresholds. A total of 1,543 to 11,438 species were detected per sample, with 59 to 10,387 species surpassing the threshold of 10 reads. The proportion of reads successfully classified at the species level ranged from 3.5% (e.g., PE, Tr1, day 2) to over 20% in some samples (e.g., wPE, Tr2, day 2, outside incubation). Despite this, a substantial number of reads remained unclassified, accounting for 50 – 95% of the total reads per sample, with the higher proportion of unclassified reads belonging to samples from the Southern Ocean (70.26% - 89.68%), reflecting either the presence of novel taxa and/or limitations in the ASW core_nt reference database used in the present study (AWS, 2024).

Nevertheless, shotgun metagenomics (Bracken) and ASV (16S rRNA gene) analyses revealed consistent ecological trends in bacterial community composition despite differences in relative abundances. In the ASV dataset, Pseudomonadota, dominated with ∼70% relative abundance, while in the Bracken dataset, they were underrepresented due to the dominance of “uncultured_bacteria.” For instance, Pseudomonadota remained prevalent across all materials and conditions in both analyses, highlighting their ecological versatility and central role in these microbial communities. Similarly, Cyanobacteria and Bacteroidota were highly detected in both analyses, with Cyanobacteria showing higher abundances on glass and water surfaces and Bacteroidota exhibiting increased prevalence on PE and wPE, reflecting their roles in oligotrophic environments and polymer degradation, respectively (Bullerjahn and Post, 2014; Fernández-Gómez et al., 2013).

Both datasets also revealed time and environment-dependent trends, with notable shifts in community composition across incubation days and between indoor and outdoor conditions. Over time, a diversification of microbial taxa was evident, particularly in the ASV data, with outdoor incubation promoting distinct changes, likely driven by sunlight and temperature. The Bracken data similarly captured these trends, though the high relative abundance of “uncultured_bacteria” obscured finer taxonomic details. Despite methodological differences, the coherence in key ecological patterns demonstrated the robustness of these microbial groups in adapting to environmental gradients and highlights the importance of refining the metagenomic classification pipeline used to improve taxonomic resolution.

#### 3.2.1 Functional profiling of polyethylene early colonising communities in South and Southern Atlantic waters

Normalised pathway abundance values (CPM) from the HUMAnN3 analysis showed that, of the total CPM, 2,626,295 CPM were classified into known pathways, while 56,825,267 CPM remained unmapped. The highest unmapped values were observed in wPE incubations during both transects at 2 and 14 days. Additionally, 11,861,586 CPM were unintegrated/unclassified, corresponding to reads mapped to individual genes but not linked to complete pathways or lacking taxonomic classification.

Further analysis of classified CPM revealed that pathway presence and contribution to community function remained relatively stable across samples (Figure 5). Mean pathway abundances concentrated at low CPM values across most samples (range of 562 – 1.3 CPM), suggesting a small number of dominant pathways and a larger proportion of rare or low-abundance pathways.

**Figure 5.**
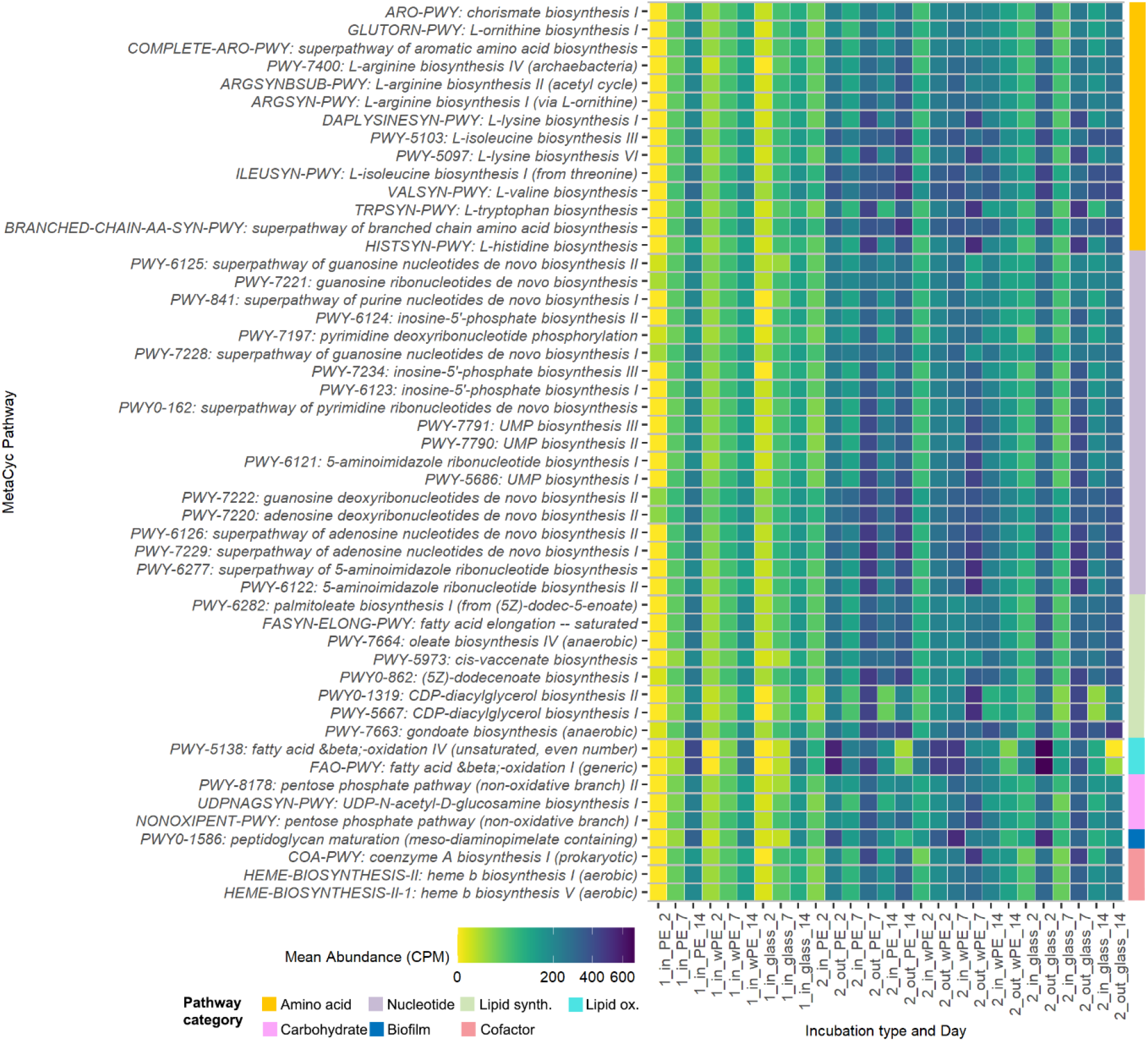
Top 50 MetaCyc microbial metabolic pathways identified across incubations. Incubation type and Day naming: 1,2 = Transect 1 or 2; in,out = incubation inside aquarium or on shipdeck; PE, wPE,glass = incubation material; 2,7,14 = biofilm harvesting day.

The 50 most abundant pathways identified across incubations were mainly dedicated to general anabolic functions, with nucleotide, amino acid, and lipid biosynthesis representing 38%, 28%, and 16% of the most abundant pathways, respectively (Fig. 5). Nonetheless, carbohydrate metabolism and lipid oxidation were also among the most abundant metabolic pathways, with fatty acid and beta oxidation as well as the pentose phosphate pathway representing 4% and 6%, respectively. Consistent with our findings, prior metaproteomic studies identified heterotrophic genera such as *Psychrobacter* and *Flavobacterium* as dominant members of the plastisphere in cold environments, with carbohydrate metabolism and oxidative phosphorylation serving as the primary metabolic pathways (Messer et al., 2024). While lipid metabolism has been proposed as a route for bacteria PE use as carbon source after initial extracellular polymer breakdown (Zadjelovic et al., 2022).

Peptidoglycan maturation was prominently identified across all systems, with a marked increase observed after 14 days in Tr1 incubations and consistently throughout Tr2 incubations. Biofilms are known to protect bacterial cells from various environmental threats, and our findings suggest a significant increase in peptidoglycan secretion (Fig. 5), in alignment with other plastisphere research (Bos et al, 2023; Monràs-Riera, Avila and Ballesté 2024). Notably, biofilm formation has been proposed to be an early bacterial response to survival challenges, particularly in *Vibrio* species, as highlighted in recent studies (Vaidya et al., 2025). Furthermore, the accessibility of PE as a source of carbon for bacteria seems to be strongly mediated by adhesion to the material’s surface (Zadjelovic et al., 2022).

Another highlight is the identification of heme b-biosynthesis among the most abundant pathways, particularly abundant by the end of the incubation period, as it has been previously proposed that iron acquisition plays a critical role in microbial colonisation of the plastisphere in marine environments, where iron is limited (Bos et al., 2023; Kim et al., 2021). Heme b is a critical component in many cellular processes vital for microbial survival and adaptability in diverse environments (Kim et al., 2021). These findings suggest that iron metabolism is central to early colonisation stages on plastic surfaces. While siderophore production and utilisation have been observed in plastisphere communities after 7 days of incubation (Bos et al., 2023), our results highlight the intracellular iron utilisation mechanisms over extracellular iron acquisition strategies, offering new insights into the metabolic priorities of plastisphere microbes.

Analysis of gene family abundances revealed a dominance of unmapped sequences, which accounted for the majority of the total abundance (∼ 5 million CPM). The highest proportion of unclassified sequences (33731 CPM, 98.3 %) was observed in the community harvested from glass after two days of incubation during Transect 1. These findings highlight a substantial fraction of unmapped and unclassified sequences, suggesting the presence of potentially novel gene families in these remote oceanic environment or incomplete annotation databases.

Among the top 100 most abundant classified gene families across incubations that could be mapped against the UniProt database, 88 % percent were closely related to families from marine microorganisms, with 26 % matched to photosynthetic functions, such as cytochrome subunits, photosystem II, and chloroplast proteins, mainly from algae, despite of eukaryotic sequences having been filtered out of the analysis, with the exception of a match to an unclassified protein from the cyanobacteria group Aphanizomenon flos-aquae, uniquely identified after 2 days on glass during Tr1. On the whole, these results confirm the observations of dominance by ASVs classified as Cyanobacteria. Other highly abundant gene families matched against Pseudomonadota, in particular of the orders Oceanospirillales, Alteromonadales, and Hypomicrobiales, commonly associated to hydrocarbon degradation, surface adhesion, biofilm formation, and nitrogen fixation respectively (Amaral-Zettler, 2022; Bos et al., 2023; Messer et al., 2024; Xu et al., 2011). Unfortunately, many of the gene families in our study were unclassified or matched against predicted proteins with unknown functions, making difficult further comparisons.

Among the highly abundant gene families, A0A2A4UQ16, uniquely identified in wPE after 7 days inside aquarium incubations during Tr2, crossed referenced (x36) to a predicted protein with unknown function of the obligate aliphatic hydrocarbon degrader, proteobacterium *Thalassolituus oleivorans* (Golyshin et al., 2013; Yakimov et al., 2004) exhibited the highest abundance (356.7 ± 40.7 CPM) across all experiments. Interestingly, although most microbes lack the specific enzymes required to initiate the breakdown of polyethylene, the degradation of pristine and weathered LDPE by *Alcanivorax sp*., a member of the Oceanospirillales order, has been recently proven (Zadjelovic et al., 2022). The proposed mechanism for PE degradation is linked to lipid metabolism, including β-oxidation enzymes such as FadB, identified in our ship deck PE incubation during Tr2 as a match to *Psychrobacter* sp.’s gene family A0A139BZ75 (fatty acid oxidation complex subunit alpha). Other gene families associated with lipid metabolism identified include A0A2A4RXG1 (Enoyl-[acyl-carrier-protein] reductase [NADH] FabI) and A0A2A4RY10 (enoyl-CoA hydratase) which are involved in the elongation cycle of fatty acids and in the complete oxidation of long-chain fatty acids, respectively. The dominance of ASVs classified as Pseudomonadota, particulary Oceanospirillales, in conjunction with gene families matches related to proposed PE degradation in aquatic environments (Zadjelovic et al., 2022) supports the hypothesis of hydrocarbon degraders enrichment in these experiments.

The identification of A0A2K9LLE0, a gene family matched to the hemolysin-coregulated protein of the n-alkane degrading *Ketobacter alkanivorans* (Kim et al., 2018) further support our hypothesis of the crucial role of iron metabolism in the early stages of PE colonisation, even more, this could be a key factor for hydrocarbon biodegradation. However, the lack of identified specific pathways or gene families, such as oxygenases, make difficult such claim.

Despite the ubiquitous and dominant identification of known hydrocarbon degraders among ASVs, the functional profiling method employed did not indicate hydrocarbon degradation as a prevalent metabolic pathway. Yet, highly abundant gene families identified included A0A2A4UQ16, A0A139BZ75, A0A2A4RY10, potentially linked to this metabolic function. Further, iron metabolism, biofilm formation, and autotrophy were recognised as highly abundant across ASVs, metabolic pathway, and gene family profiles. Unfortunately, many of the metabolic pathways and gene families in our study were unclassified or matched against predicted proteins with unknown functions. Our initial metagenomic observations warrant genome annotation and ongoing work is already examining inter-kingdom ecological roles, and future efforts will aim to expand these findings further. Nevertheless, this study already expands the understanding of plastisphere dynamics, a critical step in evaluating the ecological risks of marine plastic pollution and identifying microorganisms with promising biotechnological applications.

## Conclusion

Our results demonstrate that early colonising prokaryotic communities of drifting PE in Southern Hemisphere oceanic waters are dominated by Cyanobacteria, Pseudomonadota, and Bacteroidota, similarly to plastisphere communities in other aquatic environments. We conclude that although geography impacts abundance dynamics of these taxa, plastisphere membership does not seem to depend on location, temperature or type of type of material.

These communities showed high abundance of photosynthesis, lipid oxidation, biofilm formation, and heme b metabolic pathways and associated gene families, confirming that autotrophy, adhesion, iron metabolism, and potentially hydrocarbon degradation are key metabolic functions in the early colonisation of PE. On the whole, our results are coherent with the ecological niche that polyethylene, as a highly resistant and inert material, initially offers to early colonisers, where carbon sources are limited and specialists such as autotrophs, as well as strict hydrocarbon degraders and biofilm formers must pave the way for non-specialist heterotrophs to be able to survive. This study expands the understanding of plastisphere dynamics, particularly in Sothern Hemisphere waters and offers valuable observations for the design of plastic waste cleaning strategies.

## Acknowledgements

The authors thank Dr. Andrew Nelson for processing the raw shotgun metagenomic data on the Northumbria University high-computing cloud service. Equally, the authors thank Ms. Nicole Daniela Seiler-Kurth, for her invaluable technical support and Ms. Christine Gogel for facilitating the financial administration of this project.

We are also grateful for the logistical and technical support during expedition PS129 and PS130-1 from the entire crew of RV Polarstern: Alfred-Wegener-Institut Helmholtz-Zentrum für Polar- und Meeresforschung (2017). Polar Research and Supply Vessel POLARSTERN Operated by the Alfred-Wegener-Institute. Journal of large-scale research facilities, 3, A119. http://dx.doi.org/10.17815/jlsrf-3-163.

## Data availability

Raw amplicon and metagenomic sequencing data are openly accessible at the NCBI archive (BioProject PRJNA1195827). Sample accession numbers, sequencing processing data, as well as coordinates and other environmental data are part of the metadata available at Zenodo Data repository DOI 10.5281/zenodo.15437519.

## Author’s CRedit contributions

PCB: Conceptualisation, Methodology, Investigation, Formal analysis, Data curation, Visualisation, Writing – original draft, Project administration. GECB: Conceptualisation, Methodology, Resources. AN: Methodology. PBH: Writing – review & editing, Supervision, Funding acquisition.

## References

Aguila-Torres, P., González, M., Maldonado, J.E., Miranda, R., Zhang, L., González-Stegmaier, R., Rojas, L.A., Gaete, A., 2022. Associations between bacterial communities and microplastics from surface seawater of the Northern Patagonian area of Chile. Environmental Pollution 306, 119313. 10.1016/j.envpol.2022.119313

Amaral-Zettler, L.A., 2022. Colonization of Plastic Marine Debris. Plastics and the Ocean 301–316. 10.1002/9781119768432.CH10

Amaral-Zettler, L.A., Zettler, E.R., Mincer, T.J., 2020. Ecology of the plastisphere. Nat Rev Microbiol. 10.1038/s41579-019-0308-0

Amaral-Zettler, L.A., Zettler, E.R., Slikas, B., Boyd, G.D., Melvin, D.W., Morrall, C.E., Proskurowski, G., Mincer, T.J., 2015. The biogeography of the Plastisphere: Implications for policy. Front Ecol Environ 13, 541–546. 10.1890/150017

AWS, 2024. Amazon Web Services (AWS) [WWW Document]. AWS Public Dataset Program. URL https://benlangmead.github.io/aws-indexes/k2 (accessed 12.4.24).

Barnes, D.K.A., Galgani, F., Thompson, R.C., Barlaz, M., 2009. Accumulation and fragmentation of plastic debris in global environments. Philosophical Transactions of the Royal Society B: Biological Sciences 364, 1985–1998. 10.1098/RSTB.2008.0205

Bos, R.P., Kaul, D., Zettler, E.R., Hoffman, J.M., Dupont, C.L., Amaral-Zettler, L.A., Mincer, T.J., 2023. Plastics select for distinct early colonizing microbial populations with reproducible traits across environmental gradients. Environ Microbiol. 10.1111/1462-2920.16391

Bryant, J.A., Clemente, T.M., Viviani, D.A., Fong, A.A., Thomas, K.A., Kemp, P., Karl, D.M., White, A.E., DeLong, E.F., 2016. Diversity and Activity of Communities Inhabiting Plastic Debris in the North Pacific Gyre. mSystems 1. 10.1128/mSystems.00024-16

Bullerjahn, G.S., Post, A.F., 2014. Physiology and molecular biology of aquatic cyanobacteria. Front Microbiol. 10.3389/fmicb.2014.00359

Callahan, B.J., McMurdie, P.J., Rosen, M.J., Han, A.W., Johnson, A.J.A., Holmes, S.P., 2016. DADA2: High-resolution sample inference from Illumina amplicon data. Nat Methods 13, 581–583. 10.1038/nmeth.3869

Carrillo-Barragan, P., Fitzsimmons, C., Lloyd-Hartley, H., Tinlin-Mackenzie, A., Scott, C., Sugden, H., 2023. Fifty-year study of microplastics ingested by brachyuran and fish larvae in the central English North Sea. Environmental Pollution.

Convey, P., Peck, L.S., 2019. Antarctic environmental change and biological responses. Sci Adv 5. 10.1126/sciadv.aaz0888

Debroas, D., Mone, A., Ter Halle, A., 2017. Plastics in the North Atlantic garbage patch: A boat-microbe for hitchhikers and plastic degraders. Science of The Total Environment 599–600, 1222–1232. 10.1016/j.scitotenv.2017.05.059

Eriksen, M., Lebreton, L.C.M., Carson, H.S., Thiel, M., Moore, C.J., Borerro, J.C., Galgani, F., Ryan, P.G., Reisser, J., 2014. Plastic Pollution in the World’s Oceans: More than 5 Trillion Plastic Pieces Weighing over 250,000 Tons Afloat at Sea. PLoS One 9, e111913. 10.1371/journal.pone.0111913

Erni-Cassola, G., Wright, R.J., Gibson, M.I., Christie-Oleza, J.A., 2020. Early Colonization of Weathered Polyethylene by Distinct Bacteria in Marine Coastal Seawater. Microb Ecol 79, 517–526. 10.1007/s00248-019-01424-5

Fazey, F.M.C., Ryan, P.G., 2016. Biofouling on buoyant marine plastics: An experimental study into the effect of size on surface longevity. Environmental Pollution 210, 354–360. 10.1016/j.envpol.2016.01.026

Fernández-Gómez, B., Richter, M., Schüler, M., Pinhassi, J., Acinas, S.G., González, J.M., Pedrós-Alió, C., 2013. Ecology of marine bacteroidetes: A comparative genomics approach. ISME Journal 7, 1026–1037. 10.1038/ismej.2012.169

Franzosa, E.A., McIver, L.J., Rahnavard, G., Thompson, L.R., Schirmer, M., Weingart, G., Lipson, K.S., Knight, R., Caporaso, J.G., Segata, N., Huttenhower, C., 2018. Species-level functional profiling of metagenomes and metatranscriptomes. Nat Methods 15, 962–968. 10.1038/s41592-018-0176-y

Geyer, R., Jambeck, J.R., Law, K.L., 2017. Production, use, and fate of all plastics ever made. Sci Adv 3. 10.1126/SCIADV.1700782

Golyshin, P.N., Werner, J., Chernikova, T.N., Tran, H., Ferrer, M., Yakimov, M.M., Teeling, H., Golyshina, O. V., 2013. Genome Sequence of Thalassolituus oleivorans MIL-1 (DSM 14913 T). Genome Announc 1. 10.1128/genomeA.00141-13

Guan, X., Zhao, Z., Jiang, J., Fu, L., Liu, J., Pan, Y., Gao, S., Wang, B., Chen, Z., Wang, X., Sun, H., Jiang, B., Dong, Y., Zhou, Z., 2023. Succession and assembly mechanisms of seawater prokaryotic communities along an extremely wide salinity gradient. Environ Microbiol Rep 15, 545–556. 10.1111/1758-2229.13188

Harrison, J.P., Schratzberger, M., Sapp, M., Osborn, A.M., 2014. Rapid bacterial colonization of low-density polyethylene microplastics in coastal sediment microcosms. BMC Microbiol 14, 232. 10.1186/s12866-014-0232-4

Hoppema, M., 2023. The Expedition PS129 of the Research Vessel POLARSTERN to the Weddell Sea in 2022. 10.57738/BzPM_0776_2023

Hoppmann, M., Tippenhauer, S., Hoppema, M., 2023. Continuous thermosalinograph oceanography along RV POLARSTERN cruise track PS129 [dataset]. Bremerhaven.

Kesy, K., Oberbeckmann, S., Kreikemeyer, B., Labrenz, M., 2019. Spatial Environmental Heterogeneity Determines Young Biofilm Assemblages on Microplastics in Baltic Sea Mesocosms. Front Microbiol 10, 1665. 10.3389/FMICB.2019.01665

Kim, S., Kang, I., Lee, J.-W., Jeon, C.O., Giovannoni, S.J., Cho, J.-C., 2021. Heme auxotrophy in abundant aquatic microbial lineages. Proceedings of the National Academy of Sciences 118. 10.1073/pnas.2102750118

Kim, S.-H., Kim, J.-G., Jung, M.-Y., Kim, S.-J., Gwak, J.-H., Yu, W.-J., Roh, S.W., Kim, Y.-H., Rhee, S.-K., 2018. Ketobacter alkanivorans gen. nov., sp. nov., an n-alkane-degrading bacterium isolated from seawater. Int J Syst Evol Microbiol 68, 2258–2264. 10.1099/ijsem.0.002823

Kooi, M., Koelmans, A.A., 2019. Simplifying Microplastic via Continuous Probability Distributions for Size, Shape, and Density. Environ Sci Technol Lett 6, 551–557. 10.1021/acs.estlett.9b00379

Leistenschneider, C., Le Bohec, C., Eisen, O., Houstin, A., Neff, S., Primpke, S., Zitterbart, D.P., Burkhardt-Holm, P., Gerdts, G., 2022. No evidence of microplastic ingestion in emperor penguin chicks (Aptenodytes forsteri) from the Atka Bay colony (Dronning Maud Land, Antarctica). Science of The Total Environment 851, 158314. 10.1016/j.scitotenv.2022.158314

Leuenberger, K., Erni-Cassola, G., Leistenschneider, C., Burkhardt-Holm, P., 2024. Microplastic ingestion in five demersal, bathydemersal and bathypelagic fish species from the eastern Weddell Sea, Antarctica. Science of The Total Environment 946, 174320. 10.1016/j.scitotenv.2024.174320

McMurdie and Holmes, 2014. Shiny-phyloseq: Web Application for Interactive Microbiome Analysis with Provenance Tracking.

Messer, L.F., Lee, C.E., Wattiez, R., Matallana-Surget, S., 2024. Novel functional insights into the microbiome inhabiting marine plastic debris: critical considerations to counteract the challenges of thin biofilms using multi-omics and comparative metaproteomics. Microbiome 12, 36. 10.1186/s40168-024-01751-x

Monràs-Riera, P., Avila, C., Ballesté, E., 2024. Plastisphere in an Antarctic environment: A microcosm approach. Mar Pollut Bull 208. 10.1016/j.marpolbul.2024.116961

Oberbeckmann, S., Kreikemeyer, B., Labrenz, M., 2018. Environmental Factors Support the Formation of Specific Bacterial Assemblages on Microplastics. Front Microbiol 8. 10.3389/fmicb.2017.02709

Ogonowski, M., Motiei, A., Ininbergs, K., Hell, E., Gerdes, Z., Udekwu, K.I., Bacsik, Z., Gorokhova, E., 2018. Evidence for selective bacterial community structuring on microplastics. Environ Microbiol 20, 2796–2808. 10.1111/1462-2920.14120

Oksanen, J., 2008. Vegan: an introduction to ordination. Management 1, 1–10. https://doi.org/intro-vegan.Rnw 1260 2010-08-17 12:11:04Z jarioksa processed with vegan 1.17-6 in R version 2.12.1 (2010-12-16) on January 10, 2011.

Pörtner, H.O., 2006. Climate-dependent evolution of Antarctic ectotherms: An integrative analysis. Deep Sea Research Part II: Topical Studies in Oceanography 53, 1071–1104. 10.1016/j.dsr2.2006.02.015

Posit team, 2025. RStudio: Integrated Development Environment for R. Posit Software.

Quast, C., Pruesse, E., Yilmaz, P., Gerken, J., Schweer, T., Yarza, P., Peplies, J., Glöckner, F.O., 2012. The SILVA ribosomal RNA gene database project: improved data processing and web-based tools. Nucleic Acids Res 41, D590–D596. 10.1093/nar/gks1219

Ramachandran, A., Mclatchie, S., Walsh, D.A., 2021. A Novel Freshwater to Marine Evolutionary Transition Revealed within Methylophilaceae Bacteria from the Arctic Ocean.

Roager, L., Sonnenschein, E.C., 2019. Bacterial Candidates for Colonization and Degradation of Marine Plastic Debris. Environ Sci Technol 53, 11636–11643. 10.1021/acs.est.9b02212

Scales, B.S., Cable, R.N., Duhaime, M.B., Gerdts, G., Fischer, F., Fischer, D., Mothes, S., Hintzki, L., Moldaenke, L., Ruwe, M., Kalinowski, J., Kreikemeyer, B., Pedrotti, M.-L., Gorsky, G., Elineau, A., Labrenz, M., Oberbeckmann, S., 2021. Cross-Hemisphere Study Reveals Geographically Ubiquitous, Plastic-Specific Bacteria Emerging from the Rare and Unexplored Biosphere. mSphere 6. 10.1128/msphere.00851-20

Schleheck, D., Weiss, M., Pitluck, S., Bruce, D., Land, M.L., Han, S., Saunders, E., Tapia, R., Detter, C., Brettin, T., Han, J., Woyke, T., Goodwin, L., Pennacchio, L., Nolan, M., Cook, A.M., Kjelleberg, S., Thomas, T., 2011. Complete genome sequence of Parvibaculum lavamentivorans type strain (DS-1T). Stand Genomic Sci 5, 298–310. 10.4056/sigs.2215005

Schlundt, C., Mark Welch, J.L., Knochel, A.M., Zettler, E.R., Amaral-Zettler, L.A., 2020. Spatial structure in the “Plastisphere”: Molecular resources for imaging microscopic communities on plastic marine debris. Mol Ecol Resour 20, 620–634. 10.1111/1755-0998.13119

Sunagawa, S., Acinas, S.G., Bork, P., Bowler, C., Babin, M., Boss, E., Cochrane, G., de Vargas, C., Follows, M., Gorsky, G., Grimsley, N., Guidi, L., Hingamp, P., Iudicone, D., Jaillon, O., Kandels, S., Karp-Boss, L., Karsenti, E., Lescot, M., Not, F., Ogata, H., Pesant, S., Poulton, N., Raes, J., Sardet, C., Sieracki, M., Speich, S., Stemmann, L., Sullivan, M.B., Wincker, P., Eveillard, D., Lombard, F., 2020. Tara Oceans: towards global ocean ecosystems biology. Nat Rev Microbiol. 10.1038/s41579-020-0364-5

Sunagawa, S., Luis, †, Coelho, P., Chaffron, S., Kultima, J.R., Labadie, K., Salazar, G., Djahanschiri, B., Zeller, G., Mende, D.R., Alberti, A., Cornejo-Castillo, F.M., Costea, P.I., Cruaud, C., D’ovidio, F., Engelen, S., Ferrera, I., Gasol, J.M., Guidi, L., Hildebrand, F., Kokoszka, F., Lepoivre, C., Lima-Mendez, G., Poulain, J., Poulos, B.T., Royo-Llonch, M., Sarmento, H., Vieira-Silva, S., Dimier, C., Picheral, M., Searson, S., Kandels-Lewis, S., Pesant, S., Speich, S., Stemmann, L., Sullivan, M.B., Weissenbach, J., Wincker, P., Karsenti, E., Raes, J., Acinas, S.G., Peer Bork, †, 2015. Structure and function of the global ocean microbiome. Science (1979) 15. 10.1126/science.1261359

Telesh, I., Schubert, H., Skarlato, S., 2013. Life in the salinity gradient: Discovering mechanisms behind a new biodiversity pattern. Estuar Coast Shelf Sci. 10.1016/j.ecss.2013.10.013

Thompson, L.R., Sanders, J.G., McDonald, D., Amir, A., Ladau, J., Locey, K.J., Prill, R.J., Tripathi, A., Gibbons, S.M., Ackermann, G., Navas-Molina, J.A., Janssen, S., Kopylova, E., Vázquez-Baeza, Y., González, A., Morton, J.T., Mirarab, S., Zech Xu, Z., Jiang, L., Haroon, M.F., Kanbar, J., Zhu, Q., Jin Song, S., Kosciolek, T., Bokulich, N.A., Lefler, J., Brislawn, C.J., Humphrey, G., Owens, S.M., Hampton-Marcell, J., Berg-Lyons, D., McKenzie, V., Fierer, N., Fuhrman, J.A., Clauset, A., Stevens, R.L., Shade, A., Pollard, K.S., Goodwin, K.D., Jansson, J.K., Gilbert, J.A., Knight, R., Rivera, J.L.A., Al-Moosawi, L., Alverdy, J., Amato, K.R., Andras, J., Angenent, L.T., Antonopoulos, D.A., Apprill, A., Armitage, D., Ballantine, K., Bárta, J., Baum, J.K., Berry, A., Bhatnagar, A., Bhatnagar, M., Biddle, J.F., Bittner, L., Boldgiv, B., Bottos, E., Boyer, D.M., Braun, J., Brazelton, W., Brearley, F.Q., Campbell, A.H., Caporaso, J.G., Cardona, C., Carroll, J., Cary, S.C., Casper, B.B., Charles, T.C., Chu, H., Claar, D.C., Clark, R.G., Clayton, J.B., Clemente, J.C., Cochran, A., Coleman, M.L., Collins, G., Colwell, R.R., Contreras, M., Crary, B.B., Creer, S., Cristol, D.A., Crump, B.C., Cui, D., Daly, S.E., Davalos, L., Dawson, R.D., Defazio, J., Delsuc, F., Dionisi, H.M., Dominguez-Bello, M.G., Dowell, R., Dubinsky, E.A., Dunn, P.O., Ercolini, D., Espinoza, R.E., Ezenwa, V., Fenner, N., Findlay, H.S., Fleming, I.D., Fogliano, V., Forsman, A., Freeman, C., Friedman, E.S., Galindo, G., Garcia, L., Garcia-Amado, M.A., Garshelis, D., Gasser, R.B., Gerdts, G., Gibson, M.K., Gifford, I., Gill, R.T., Giray, T., Gittel, A., Golyshin, P., Gong, D., Grossart, H.-P., Guyton, K., Haig, S.-J., Hale, V., Hall, R.S., Hallam, S.J., Handley, K.M., Hasan, N.A., Haydon, S.R., Hickman, J.E., Hidalgo, G., Hofmockel, K.S., Hooker, J., Hulth, S., Hultman, J., Hyde, E., Ibáñez-Álamo, J.D., Jastrow, J.D., Jex, A.R., Johnson, L.S., Johnston, E.R., Joseph, S., Jurburg, S.D., Jurelevicius, D., Karlsson, A., Karlsson, R., Kauppinen, S., Kellogg, C.T.E., Kennedy, S.J., Kerkhof, L.J., King, G.M., Kling, G.W., Koehler, A. V., Krezalek, M., Kueneman, J., Lamendella, R., Landon, E.M., Lane-deGraaf, K., LaRoche, J., Larsen, P., Laverock, B., Lax, S., Lentino, M., Levin, I.I., Liancourt, P., Liang, W., Linz, A.M., Lipson, D.A., Liu, Y., Lladser, M.E., Lozada, M., Spirito, C.M., MacCormack, W.P., MacRae-Crerar, A., Magris, M., Martín-Platero, A.M., Martín-Vivaldi, M., Martínez, L.M., Martínez-Bueno, M., Marzinelli, E.M., Mason, O.U., Mayer, G.D., McDevitt-Irwin, J.M., McDonald, J.E., McGuire, K.L., McMahon, K.D., McMinds, R., Medina, M., Mendelson, J.R., Metcalf, J.L., Meyer, F., Michelangeli, F., Miller, K., Mills, D.A., Minich, J., Mocali, S., Moitinho-Silva, L., Moore, A., Morgan-Kiss, R.M., Munroe, P., Myrold, D., Neufeld, J.D., Ni, Y., Nicol, G.W., Nielsen, S., Nissimov, J.I., Niu, K., Nolan, M.J., Noyce, K., O’Brien, S.L., Okamoto, N., Orlando, L., Castellano, Y.O., Osuolale, O., Oswald, W., Parnell, J., Peralta-Sánchez, J.M., Petraitis, P., Pfister, C., Pilon-Smits, E., Piombino, P., Pointing, S.B., Pollock, F.J., Potter, C., Prithiviraj, B., Quince, C., Rani, A., Ranjan, R., Rao, S., Rees, A.P., Richardson, M., Riebesell, U., Robinson, C., Rockne, K.J., Rodriguezl, S.M., Rohwer, F., Roundstone, W., Safran, R.J., Sangwan, N., Sanz, V., Schrenk, M., Schrenzel, M.D., Scott, N.M., Seger, R.L., Seguin-Orlando, A., Seldin, L., Seyler, L.M., Shakhsheer, B., Sheets, G.M., Shen, C., Shi, Y., Shin, H., Shogan, B.D., Shutler, D., Siegel, J., Simmons, S., Sjöling, S., Smith, D.P., Soler, J.J., Sperling, M., Steinberg, P.D., Stephens, B., Stevens, M.A., Taghavi, S., Tai, V., Tait, K., Tan, C.L., Taş, N., Taylor, D.L., Thomas, T., Timling, I., Turner, B.L., Urich, T., Ursell, L.K., van der Lelie, D., Van Treuren, W., van Zwieten, L., Vargas-Robles, D., Thurber, R.V., Vitaglione, P., Walker, D.A., Walters, W.A., Wang, S., Wang, T., Weaver, T., Webster, N.S., Wehrle, B., Weisenhorn, P., Weiss, S., Werner, J.J., West, K., Whitehead, A., Whitehead, S.R., Whittingham, L.A., Willerslev, E., Williams, A.E., Wood, S.A., Woodhams, D.C., Yang, Y., Zaneveld, J., Zarraonaindia, I., Zhang, Q., Zhao, H., 2017. A communal catalogue reveals Earth’s multiscale microbial diversity. Nature 551, 457–463. 10.1038/nature24621

Thompson, R.C., Swan, S.H., Moore, C.J., vom Saal, F.S., 2009. Our plastic age. Philosophical Transactions of the Royal Society B: Biological Sciences 364, 1973–1976. 10.1098/rstb.2009.0054

Vaidya, S., Saha, D., Rode, D.K.H., Torrens, G., Hansen, M.F., Singh, P.K., Jelli, E., Nosho, K., Jeckel, H., Göttig, S., Cava, F., Drescher, K., 2025. Bacteria use exogenous peptidoglycan as a danger signal to trigger biofilm formation. Nat Microbiol 10, 144–157. 10.1038/s41564-024-01886-5

Wilkie Johnston, L., Bergami, E., Rowlands, E., Manno, C., 2023. Organic or junk food? Microplastic contamination in Antarctic krill and salps. R Soc Open Sci 10. 10.1098/rsos.221421

Wright, R.J., Erni-Cassola, G., Zadjelovic, V., Latva, M., Christie-Oleza, J.A., 2020. Marine Plastic Debris: A New Surface for Microbial Colonization. Environ Sci Technol 54, 11657–11672. 10.1021/acs.est.0c02305

Xu, X.-W., Huo, Y.-Y., Wang, C.-S., Oren, A., Cui, H.-L., Vedler, E., Wu, M., 2011. Pelagibacterium halotolerans gen. nov., sp. nov. and Pelagibacterium luteolum sp. nov., novel members of the family Hyphomicrobiaceae. Int J Syst Evol Microbiol 61, 1817–1822. 10.1099/ijs.0.023325-0

Yakimov, M.M., Giuliano, L., Denaro, R., Crisafi, E., Chernikova, T.N., Abraham, W.-R., Luensdorf, H., Timmis, K.N., Golyshin, P.N., 2004. Thalassolituus oleivorans gen. nov., sp. nov., a novel marine bacterium that obligately utilizes hydrocarbons. Int J Syst Evol Microbiol 54, 141–148. 10.1099/ijs.0.02424-0

Zadjelovic, V., Erni-Cassola, G., Obrador-Viel, T., Lester, D., Eley, Y., Gibson, M.I., Dorador, C., Golyshin, P.N., Black, S., Wellington, E.M.H., Christie-Oleza, J.A., 2022. A mechanistic understanding of polyethylene biodegradation by the marine bacterium Alcanivorax. J Hazard Mater 436. 10.1016/j.jhazmat.2022.129278

Zeghal, E., Vaksmaa, A., van Bleijswijk, J., Niemann, H., 2024. Environmental factors control microbial colonization of plastics in the North Sea. Mar Pollut Bull 208. 10.1016/j.marpolbul.2024.116964

Zettler, E.R., Mincer, T.J., Amaral-Zettler, L.A., 2013. Life in the “Plastisphere”: Microbial Communities on Plastic Marine Debris. 10.1021/es401288x

