## SupplementaryTable1 for "Microbial Colonisation of Polyethylene in Offshore Marine Environments: Insights from the Southern and South Atlantic Oceans"

### Supplementary Table 1

| Expedition | Sampling Station* | Sample ID* | Col. date | Location | Coordinates DD | | T (°C) | Salinity (PSS-78) | Mol. bio. analyses | Reference(s) |
|---|---|---|---|---|---|---|---|---|---|---|
| | | | | | Latitude | Longitude | | | | |
| PS129 | Tr1_d2 | Tr1_d2_in | 13.04.2022 | SO | -66.11 | -31.84 | -1.74 | 34.8 | V4 16S RNA | (Hoppema, 2023; Hoppmann et al., 2023a), BioProject PRJNA1195827 |
| PS129 | Tr1_d7 | Tr1_d7_in | 18.04.2022 | SO | -64.54 | -44.86 | 1 | 33.9 | V4 16S RNA | |
| PS129 | Tr1_14 | Tr1_14_in | 25.04.2022 | SO | -58.25 | -60.33 | 1.07 | 34.7 | V4 16S RNA | |
| PS130/1 | Tr2_d2 | Tr2_d2_in | 04.05.2022 | SAO | -39.63 | -48.97 | 19.35 | 36.2 | V4 16S RNA | |
| PS130/1 | Tr2_d2 | Tr2_d2_out | 04.05.2022 | SAO | -39.63 | -48.97 | 19.35 | 36.2 | V4 16S RNA | |
| PS130/1 | Tr2_d7 | Tr2_d7_in | 09.05.2022 | SAO | -20.52 | -33.93 | 23.53 | 40.6 | V4 16S RNA | |
| PS130/1 | Tr2_d7 | Tr2_d7_out | 09.05.2022 | SAO | -20.52 | -33.93 | 23.53 | 40.6 | V4 16S RNA | |
| PS130/1 | Tr2_d14 | Tr2_d14_in | 16.05.2022 | NAO | 7.66 | -23.49 | 24.11 | 39.1 | V4 16S RNA | |
| PS130/1 | Tr2_d14 | Tr2_d14_out | 16.05.2022 | NAO | 7.66 | -23.49 | 24.11 | 39.1 | V4 16S RNA | |
| Sorcerer II | G (d3) | 18-Drifterexpttime3punches | 22.03.2017 | NAO | 25.86 | -68.99 | 26.2 | na | WGS | (Bos et al., 2023) |
| Sorcerer II | L (d5) | 49-Drifterexpttime5punches | 24.03.2017 | NAO | 22.56 | -63.73 | 27.5 | na | WGS | |
| Sorcerer II | L (d7) | 53-Drifterexpttime7punches | 26.03.2017 | NAO | 22.56 | -63.73 | 27.5 | na | WGS | |
| ASRS | ASRS | UV-weathered PE pellets | Jan-Mar. 2022 | SO | -62.67 | -60.39 | 0 ± 2 | na | 16S RNA | (Monràs-Riera et al., 2024) |
| Tara | TARA_076 | TARA_B100000513 | 16.10.2010 | SAO | -20.94 | -35.18 | 23.3 | 37.1 | WGS | (Sunagawa et al., 2015) |
| Tara | TARA_078 | TARA_B100000524 | 04.11.2010 | SAO | -30.14 | -43.29 | 19.9 | 36.3 | WGS | |
| Tara | TARA_084 | TARA_B100000780 | 03.01.2011 | SO | -60.23 | -60.65 | 1.8 | 33.7 | WGS | |
| Tara | TARA_085 | TARA_B100000787 | 06.01.2011 | SO | -62.04 | -49.53 | 0.7 | 34.4 | WGS | |

- *Sampling station and sample ID as recorded in the original reference. For PS129 and PS130/1, Tr = Transect, d = days incubated, “in”/”out” = location of incubations, inside aquarium or outside on ship deck.
- Col. date = collection date
- SO = Southern Ocean; SAO = South Atlantic Ocean; NOA = North Atlantic Ocean
- DD = Decimal Notation
- PSS-78 = Practical Salinity Scale 1978 (UNESCO, 1981)
- ASRS = Antarctic Spanish Research Station (ASRS) in Livingston Island (South Shetlands, Antarctica)

PS129-130: ocean temperature and salinity data were recorded at a depth of 11m along the cruise track. Raw data acquired by two SBE21 thermosalinographs and two auxiliary SBE38 temperature sensors (Sea-Bird Scientific, USA) installed in an underway seawater flow-through system on board RV Polarstern were processed to yield a calibrated and validated data set of temperature and salinity along the cruise track (Hoppmann et al., 2023b)

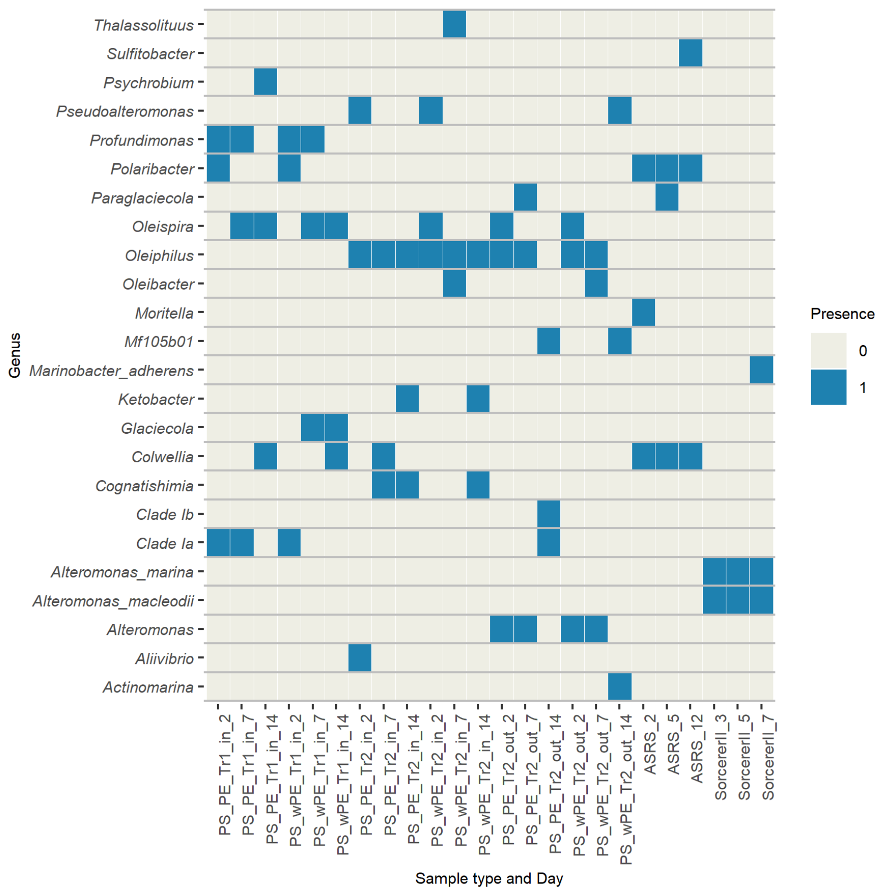


**Figure S1.** Heatmap comparing the three most abundant genra in the present Polarstern incubations versus those in the ASRS (Monràs-Riera et al., 2024) and Sorcerer II (Bos et al., 2023) studies.

Bos, R. P., Kaul, D., Zettler, E. R., Hoffman, J. M., Dupont, C. L., Amaral-Zettler, L. A., & Mincer, T. J. (2023). Plastics select for distinct early colonizing microbial populations with reproducible traits across environmental gradients. *Environmental Microbiology*. https://doi.org/10.1111/1462-2920.16391

Hoppema, M. (2023). *The Expedition PS129 of the Research Vessel POLARSTERN to the Weddell Sea in 2022*. https://doi.org/https://doi.org/10.57738/BzPM_0776_2023

Hoppmann, M., Tippenhauer, S., & Hoppema, M. (2023a). *Continuous thermosalinograph oceanography along RV POLARSTERN cruise track PS129*. PANGAEA. https://doi.org/10.1594/PANGAEA.955769

Hoppmann, M., Tippenhauer, S., & Hoppema, M. (2023b). *Continuous thermosalinograph oceanography along RV POLARSTERN cruise track PS129 [dataset]*.

Monràs-Riera, P., Avila, C., & Ballesté, E. (2024). Plastisphere in an Antarctic environment: A microcosm approach. *Marine Pollution Bulletin*, *208*. https://doi.org/10.1016/j.marpolbul.2024.116961

Sunagawa, S., Luis, †, Coelho, P., Chaffron, S., Kultima, J. R., Labadie, K., Salazar, G., Djahanschiri, B., Zeller, G., Mende, D. R., Alberti, A., Cornejo-Castillo, F. M., Costea, P. I., Cruaud, C., D'ovidio, F., Engelen, S., Ferrera, I., Gasol, J. M., Guidi, L., ... Peer Bork, †. (2015). Structure and function of the global ocean microbiome. *Science*, *15*(348). https://doi.org/DOI: 10.1126/science.1261359

UNESCO. (1981). *Background papers and supporting data on the Practical Salinity Scale 1978*.
