## SupplementaryTable2 for "Microbial Colonisation of Polyethylene in Offshore Marine Environments: Insights from the Southern and South Atlantic Oceans"

| Year | Reference | Sequencing Method+ | Location | Primers | Reference |
|---|---|---|---|---|---|
| 2013 | Zettler et al. (2013) | V4-V6 16S / V7 18S rRNA | North Atlantic Ocean | 518f / 1046r | E. R. Zettler, T. J. Mincer, L. A. Amaral-Zettler, Life in the "plastisphere": microbial communities on plastic marine debris. *Environmental Science and Technology* , **47**, 7137-7146 (2013). |
| 2014 | Harrison et al. (2014) | 16S rRNA | North Sea | 27f / 1492r | J. P. Harrison, M. Schratzberger, M. Sapp, A. M. Osborn, Rapid bacterial colonization of low-density polyethylene microplastics in coastal sediment microcosms. *BMC Microbiology* , ***14*** , 1-15 (2014). |
| 2014 | Oberbeckmann et al. (2014) | 16S rRNA | North Sea | 341f / 534r | S. Oberbeckmann, M. G. Loeder, G. Gerdts, A. M. Osborn, Spatial and seasonal variation in diversity and structure of microbial biofilms on marine plastics in Northern European waters. *FEMS Microbiology Ecology* , **90**, 478-492 (2014). |
| 2015 | Amaral-Zettler et al. (2015) | V6 16S rRNA | North Atlantic and Pacific Gyres | 967f / 1064r | L. A. Amaral-Zettler, E. R. Zettler, B. Slikas, G. D. Boyd, D. W. Melvin, C. E. Morrall, et al., The biogeography of the Plastisphere: implications for policy. *Frontiers in Ecology and the Environment* , **13**, 541-546 (2015). |
| 2015 | De Tender et al. (2015) | V3-V4 16S rRNA | North Sea | 341f / 785r | C. A. De Tender, L. I. Devriese, A. Haegeman, S. Maes, T. Ruttink, P. Dawyndt, Bacterial community profiling of plastic litter in the Belgian part of the North Sea. *Environmental Science and Technology* , ***49*** , 9629-9638(2015). |
| 2016 | Bryant et al. (2016) | WGS | North Pacific Subtropical Gyre | NA | J. A. Bryant, T. M. Clemente, D. A. Viviani, A. A. Fong, K. A. Thomas, P. Kemp, et al., Diversity and activity of communities inhabiting plastic debris in the North Pacific Gyre. *MSystems* , **1**, e00024-16 (2016). |
| 2016 | Kesy et al. (2016) | 16S rRNA | Baltic Sea | Com1f / Com2-Phr | K. Kesy, S. Oberbeckmann, F. Müller, M. Labrenz, Polystyrene influences bacterial assemblages in *Arenicola marina* -populated aquatic environments in vitro. *Environmental Pollution* , ***219*** , 219-227 (2016). |
| 2016 | Kirstein et al. (2016) | V3-V4 16S rRNA | North Sea | Various | I. V. Kirstein, S. Kirmizi, A. Wichels, A. Garin-Fernandez, R. Erler, M. Löder, G. Gerdts, Dangerous hitchhikers? Evidence for potentially pathogenic *Vibrio* spp. on microplastic particles. *Marine Environmental Research* , ***120*** , 1-8 (2016). |
| 2016 | McCormick et al. (2016) | V4 16S rRNA | Chicago River | 515f / 806r | A. R. McCormick, T. J. Hoellein, M. G. London, J. Hittie, J. W. Scott, J. J. Kelly, Microplastic in surface waters of urban rivers: concentration, sources, and associated bacterial assemblages. *Ecosphere* , ***7*** , e01556 (2016). |
| 2016 | Oberbeckmann et al. (2016) | V4 16S / V9 18S rRNA | North Sea | 515f / 806r; 1391f / 1795r | S. Oberbeckmann, A. M. Osborn, M. B. Duhaime, Microbes on a bottle: substrate, season and geography influence community composition of microbes colonizing marine plastic debris. *PLoS One* , **11,** (2016). |
| 2017 | De Tender et al. (2017) | V3-V4 16S rRNA / ITS2 | North Sea | 341f / 785r; fITS7b / ITS4NGSr | C. De Tender, L. I. Devriese, A. Haegeman, S. Maes, J. Vangeyte, A. Cattrijsse, et al., Temporal dynamics of bacterial and fungal colonization on plastic debris in the North Sea. *Environmental Science and Technology* , **51**, 7350-7360 (2017). |
| 2017 | Debroas et al. (2017) | V4 16S / V7 18S rRNA | North Atlantic Gyre | 515f / 806r; 960f / NSR1438 | D. Debroas, A. Mone, A.Ter Halle, Plastics in the North Atlantic garbage patch: a boat-microbe for hitchhikers and plastic degraders. *Science of the total environment* , ***599*** , 1222-1232 (2017) |
| 2017 | Hoellein et al. (2017) | V4 16S rRNA | Dupage River | 515f / 806r | T. J. Hoellein, A. R. McCormick, J. Hittie, M. G. London, J. W. Scott, J. J. Kelly, Longitudinal patterns of microplastic concentration and bacterial assemblages in surface and benthic habitats of an urban river. *Freshwater Science* , ***36*** , 491-507 (2017). |
| 2017 | Kettner et al. (2017) | V4 18S rRNA | Baltic Sea | 565f / 981r | M. T. Kettner, K. Rojas-Jimenez, S. Oberbeckmann, M. Labrenz, H. P. Grossart, Microplastics alter composition of fungal communities in aquatic ecosystems. *Environmental Microbiology* , ***19*** , 4447-4459 (2017). |
| 2017 | Syranidou et al. (2017) | 16S rRNA / 23S rRNA | Laboratory | ITSF / ITSReub | E. Syranidou, K. Karkanorachaki, F. Amorotti, M. Franchini, E. Repouskou, M. Kaliva, N. Kalogerakis, Biodegradation of weathered polystyrene films in seawater microcosms. *Scientific Reports* , ***7*** , 1-12 (2017). |
| 2018 | Dussud et al. (2018a) | 16S rRNA | Mediterranean Sea | 515f-y / 926r | C. Dussud, A. L. Meistertzheim, P. Conan, M. Pujo-Pay, M. George, P. Fabre, et al., Evidence of niche partitioning among bacteria living on plastics, organic particles and surrounding seawaters. *Environmental Pollution* , ***236*** **,** 807-816 (2018) |
| 2018 | Dussud et al. (2018b) | 16S rRNA | Mediterranean Sea | 515f-y / 926r | C. Dussud, C. Hudec, M. George, P. Fabre, P. Higgs, S. Bruzaud, et al., Colonization of non-biodegradable and biodegradable plastics by marine microorganisms. *Frontiers in Microbiology* , ***9*** , 1571 (2018). |
| 2018 | Frere et al. (2018) | V4-V5 16S rRNA | Bay of Breast | 518f / 926r | L. Frère, L. Maignien, M. Chalopin, A. Huvet, E. Rinnert, H. Morrison, et al., Microplastic bacterial communities in the Bay of Brest: Influence of polymer type and size. *Environmental Pollution* , ***242*** , 614-625 (2018). |
| 2018 | Jiang et al. (2018) | V3-V4 16S rRNA | East China Sea | 319f / 806r | P. Jiang, S. Zhao, L. Zhu, D. Li, Microplastic-associated bacterial assemblages in the intertidal zone of the Yangtze Estuary. *Science of the Total Environment* , ***624*** , 48-54 (2018). |
| 2018 | Kirstein et al. (2018) | V3-V4 16S rRNA / V4 18S rRNA | North Sea | 341f / 785r ; Eu565F / EU981R | I. V. Kirstein, A. Wichels, G. Krohne, G. Gerdts, Mature biofilm communities on synthetic polymers in seawater-Specific or general? *Marine Environmental Research* , ***142*** , 147-154 (2018). |
| 2018 | Oberbeckmann et al. (2018) | V4 16S rRNA | Baltic Sea | 515f / 806r | S. Oberbeckmann, B. Kreikemeyer, M. Labrenz, Environmental factors support the formation of specific bacterial assemblages on microplastics. *Frontiers in Microbiology* , **8**, 2709 (2018). |
| 2018 | Ognowski et al. (2018) | V6 16S rRNA | Baltic Sea | 534f / 783r | M. Ogonowski, A. Motiei, K. Ininbergs, E. Hell, Z. Gerdes, K. I. Udekwu, et al., Evidence for selective bacterial community structuring on microplastics. *Environmental Microbiology* , **20**, 2796-2808 (2018). |
| 2018 | Pollet et al. (2018) | 16S rRNA | Toulon Bay, France, Mediterranean Sea | 515f-Y/926r | T. Pollet, L. Berdjeb, C. Garnier, G. Durrieu, C. Le Poupon, B. Misson, J. F. Briand, Prokaryotic community successions and interactions in marine biofilms: the key role of Flavobacteriia. *FEMS Microbiology Ecology* , **94**, fiy083 (2018). |
| 2018 | Woodall et al. (2018) | V4 16S rRNA | equatorial Atlantic Ocean | 515f / 806r | L. C. Woodall, A. Sanchez-Vidal, M. Canals, G. L. Paterson, R. Coppock, V. Sleight, R. C. Thompson, The deep sea is a major sink for microplastic debris. *Royal Society Open Science* , ***1*** , 140317 (2014). |
| 2019 | Curren and Leong (2019) | V3-V4 16S rRNA | Singapore Strait, Singapore | 515f-806r | E. Curren, S. C. Y. Leong, Profiles of bacterial assemblages from microplastics of tropical coastal environments. *Science of the Total Environment* , **655**, 313-320 (2019). |
| 2019 | Delacuvellerie et al. (2019) | V3-V4 16S rRNA | Mediterranean Sea | 515f-806r | A. Delacuvellerie, V. Cyriaque, S. Gobert, S. Benali, R. Wattiez, The plastisphere in marine ecosystem hosts potential specific microbial degraders including *Alcanivorax borkumensis* as a key player for the low-density polyethylene degradation. *Journal of Hazardous Materials* , ***380*** , 120899 (2019). |
| 2019 | Erni-Cassola et al. (2019) | V4-V5 16S rRNA | Mediterranean Sea | 515f-y / 926r | G. Erni-Cassola, R. J. Wright, M. I. Gibson, J. A. Christie-Oleza, Early colonization of weathered polyethylene by distinct bacteria in marine coastal seawater. *Microbial Ecology* , **79**, 517-526 (2020). |
| 2019 | Kesy et al. (2019) | V4 16S rRNA | Baltic Sea | 515f / 806r | K. Kesy, S. Oberbeckmann, B. Kreikemeyer, M. Labrenz, M. Spatial environmental heterogeneity determines young biofilm assemblages on microplastics in Baltic Sea mesocosms. *Frontiers in Microbiology* , **10**, 1665 (2019). |
| 2019 | Kettner et al. (2019) | V4 18S rRNA | Northeast Germany | Eu565F / Eu981R | M. T. Kettner, S. Oberbeckmann, M. Labrenz, H. P. Grossart, The eukaryotic life on microplastics in brackish ecosystems. *Frontiers in Microbiology* , ***10*** , 538 (2019). |
| 2019 | Kirstein et al. (2019) | V3-V4 16S rRNA | North Sea | 341f / 785r | I. V. Kirstein, A. Wichels, E. Gullans, G. Krohne, G. Gerdts, The plastisphere–uncovering tightly attached plastic "specific" microorganisms. *PLoS One* , ***14*** , e0215859 (2019). |
| 2019 | Pinnel and Turner (2019) | WGS | Laguna Madre Lagoon TX, USA | NA | L. J. Pinnell, J. W. Turner, Shotgun metagenomics reveals the benthic microbial community response to plastic and bioplastic in a coastal marine environment. *Frontiers in Microbiology* , ***10*** , 1252 (2019). |
| 2019 | Pinto et al. (2019) | V3-V4 16S rRNA | Northern Adriatic Sea | 341f / 802r | M. Pinto, T. M., Langer, T. Hüffer, T. Hofmann, G. J., Herndl, G. J., The composition of bacterial communities associated with plastic biofilms differs between different polymers and stages of biofilm succession. *PLoS One* , **14**, (2019). |
| 2019 | Syranidou et al. (2019) | V4 16S rRNA | Laboratory | 515f / 806r | E. Syranidou, K. Karkanorachaki, F. Amorotti, A. Avgeropoulos, B. Kolvenbach, N. Y. Zhou, N. Kalogerakis, Biodegradation of mixture of plastic films by tailored marine consortia. *Journal of Hazardous Materials* , ***375*** , 33-42 (2019). |
| 2019 | Tagg et al. (2019) | V4 16S rRNA | Baltic Sea | 515f / 806r | A. S. Tagg, J. A. I. do Sul, Is this your glitter? An overlooked but potentially environmentally-valuable microplastic. *Marine Pollution Bulletin* , ***146*** , 50-53 (2019). |
| 2019 | Wu et al. (2019) | V3-V4 16S rRNA; WGS | Haihe River | 338f / 806r | X. Wu, J. Pan, M. Li, Y. Li, M. Bartlam, Y. Wang, Selective enrichment of bacterial pathogens by microplastic biofilm. *Water Research* , ***165*** , 114979 (2019). |
| 2019 | Xu et al. (2019) | V3-V4 16S rRNA | Yellow Sea | 341f / 785r | X. Xu, S. Wang, F. Gao, J. Li, L. Zheng, C. Sun, et al., Marine microplastic-associated bacterial community succession in response to geography, exposure time, and plastic type in China's coastal seawaters. *Marine Pollution Bulletin* , **145**, 278-286 (2019). |
| 2020 | Dudek et al. (2020) | V4-V5 16S rRNA / V4 18S rRNA | Caribbean Sea | 515f / 926r; eukv4F / eukv4R | K. L. Dudek, B. N. Cruz, B. Polidoro, B., S. Neuer, Microbial colonization of microplastics in the Caribbean Sea. *Limnology and Oceanography Letters* , **5**, 5-17 (2020). |
| 2020 | Lacerda et al. (2020) * | ITS2, V4 and V9 18S rRNA | South Atlantic Ocean | ITS1f/ITS4;<br>TAReuk454FWD1/TAReukREV3;<br>1391f/Euk B | Lacerda A.L.D.F.; Proietti M.C.; Secchi E.R.; Taylor J.D., Diverse groups of fungi are associated with plastics in the surface waters of the Western South Atlantic and the Antarctic Peninsula. Molecular Ecology, 29, 1903-1918 (2020) |
| 2020 | Muthukrishnan et al. (2020) | V4 16S rRNA | Marina Shangri La, Muscat Oman | 341f / 805r | R. M. Abed, T. Muthukrishnan, M. Al Khaburi, F. Al-Senafi, A. Munam, H. Mahmoud, Degradability and biofouling of oxo-biodegradable polyethylene in the planktonic and benthic zones of the Arabian Gulf. *Marine Pollution Bulletin* , ***150*** , 110639 (2020). |
| 2020 | Schlundt et al. (2020) | 16S rRNA | Vineyard Sound, True Blue Bay, North Sea | Various | C. Schlundt, J. L. Mark Welch, A. M. Knochel, E. R. Zettler, L. A. Amaral-Zettler, Spatial structure in the "Plastisphere": Molecular resources for imaging microscopic communities on plastic marine debris. Molecular Ecology Resources, 20, 620-634 (2020). |
| 2021 | Agostini et al. (2021) * | V4 16S rRNA | Brazilian continental shelf ( NADW/LCDW), South Atlantic | 515f / 806r | Agostini L.; Moreira J.C.F.; Bendia A.G.; Kmit M.C.P.; Waters L.G.; Santana M.F.M.; Sumida P.Y.G.; Turra A.; Pellizari V.H., Deep-sea plastisphere: Long-term colonization by plastic-associated bacterial and archaeal communities in the Southwest Atlantic Ocean, Science of the Total Environment, 793, 148335 (2021) |
| 2021 | Bhagwat et al. (2021) | WGS | Lake Macquarie, Australia | NA | G. Bhagwat, Q. Zhu, W. O'Connor, S. Subashchandrabose, I. Grainge, R. Knight, et al., Exploring the composition and functions of plastic microbiome using whole-genome sequencing. *Environmental Science and Technology* , **55**, 4899-4913 (2021). |
| 2021 | Cappello et al., 2021 * | Total RNA-> cDNA-> 16S rRNA | King George Island, Antarctica, Southern Ocean | 530f/192r;<br>pGEM T-easy vector M13f/M13r | appello, S., Caruso, G., Bergami, E., Macrì, A., Venuti, V., Majolino, D., & Corsi, I. New insights into the structure and function of the prokaryotic communities colonizing plastic debris collected in King George Island (Antarctica): Preliminary observations from two plastic fragments. Journal of Hazardous Materials, 414. (2021 |
| 2021 | Catão et al. (2021) | V4 16S rRNA | Mediterranean Sea, North Atlantic Ocean, Indian Ocean | 515F-Y/926R | Catão C. P. E.; Pollet T.; Garnier C.; Barry-Martinet R.; Rehel K.; Linossier I.; Tunin-Ley A.; Turquet J.; Briand J.-F., Temperate and tropical coastal waters share relatively similar microbial biofilm communities while free-living or particle-attached communities are distinct, Molecular Ecology, 30, 2891-2904, (2021) |
| 2021 | Näkki et al. (2021) | V3-V4 16S rRNA | Helsinki Harbour, Finland | 341F / 785R | Näkki P.; Eronen-Rasimus E.; Kaartokallio H.; Kankaanpää H.; Setälä O.; Vahtera E.; Lehtiniemi M., Polycyclic aromatic hydrocarbon sorption and bacterial community composition of biodegradable and conventional plastics incubated in coastal sediments, Science of the Total Environment, 755, 143088 (2021) |
| 2021 | Oberbeckmann et al. (2021) | WGS | Baltic Sea | NA | S. Oberbeckmann, D. Bartosik, S. Huang, J. Werner, C. Hirschfeld, D. Wibberg, et al., Genomic and proteomic profiles of biofilms on microplastics are decoupled from artificial surface properties. *Environmental Microbiology* , (2021). |
| 2022 | Caroppo et al., 2022 * | NA (Imaging and ezymatic activity assays) | Ross Sea, Antarctica, Southern Ocean | NA | Caroppo, C., Azzaro, M., Dell'Acqua, O., Azzaro, F., Maimone, G., Rappazzo, A. C., Raffa, F., & Caruso, G. (2022). Microbial Biofilms Colonizing Plastic Substrates in the Ross Sea (Antarctica). Journal of Marine Science and Engineering, 10,11, (2022). |
| 2022 | Lacerda et al. (2022) * | V4 16S RNA; V4 and V9 18S rRNA | Western South Atlantic Ocean | 515f/806r; TAReuk454/TAReukRev3;<br>1391f /EukB | Lacerda A.L.D.F.; Taylor J.D.; Rodrigues L.D.S.; Kessler F.; Secchi E.; Proietti M.C., Floating plastics and their associated biota in the Western South Atlantic, Floating plastics and their associated biota in the Western South Atlantic, Science of the Total Environment, 805, 150186, (2022) |
| 2022 | Kelly et al. (2022) | V4 16S rRNA | Deep-sea North-East Atlantic Ocean | Schloss Wet Lab primer set | Kelly M.R.; Whitworth P.; Jamieson A.; Burgess J.G., Bacterial colonisation of plastic in the Rockall Trough, North-East Atlantic: An improved understanding of the deep-sea plastisphere, Environmental Pollution, 305, 119314 (2022) |
| 2023 | Bos et al. (2023) | WGS | North Atlantic Ocean | NA | Bos RP, Kaul D, Zettler ER, Hoffman JM, Dupont CL, Amaral-Zettler LA, Mincer TJ., Plastics select for distinct early colonizing microbial populations with reproducible traits across environmental gradients, Environmental Microbiology, 12, 2761-2775 (2023) |
| 2023 | Caruso et al., 2023 * | Carbon substrate utilisation | Ross Sea, Antarctica, Southern Ocean | NA | Caruso, G., Maimone, G., Rappazzo, A. C., Dell'Acqua, O., Laganà, P., & Azzaro, M. Microbial Biofilm Colonizing Plastic Substrates in the Ross Sea (Antarctica): First Overview of Community-Level Physiological Profiles. Journal of Marine Science and Engineering, 11(7), 1317. (2023) |
| 2023 | Mincer et al. (2023) | WGS | North Atlantic Ocean | hsp60 gene sequence (*Vibrio* sp.) | Mincer T.J.; Bos R.P.; Zettler E.R.; Zhao S.; Asbun A.A.; Orsi W.D.; Guzzetta V.S.; Amaral-Zettler L.A., Sargasso Sea Vibrio bacteria: Underexplored potential pathovars in a perturbed habitat, Water Research, 242, 120033, (2023) |
| 2024 | Monràs-Riera et al. (2024) * | V4 16S rRNA; qPCR 16S 341F and 534R | South Shetlands, Antarctica, Southern Ocean | 515f / 806r; qPCR 16S 341F and 534R | Monràs-Riera, P., Avila, C., & Ballesté, E. Plastisphere in an Antarctic environment: A microcosm approach. Marine Pollution Bulletin, 208, (2024) |
| 2024 | Papale et al., 2024 * | V3-V4 16S rRNA | Terra Nova Bay, Antarctica, Southern Ocean | 515f / 806r | Papale, M., Fazi, S., Severini, M., Scarinci, R., Dell'Acqua, O., Azzaro, M., Venuti, V., Fazio, B., Fazio, E., Crupi, V., Irrera, A., Rizzo, C., Giudice, A. Lo, & Caruso, G. Structural properties and microbial diversity of the biofilm colonizing plastic substrates in Terra Nova Bay (Antarctica). Science of the Total Environment, 943. (202 |
| 2024 | Tigreros-Benavides et al. (2024) | V3-V4 16S rRNA | Caribean Sea | Bakt_341F / Bakt_805R | Tigreros-Benavides P.; Garzón-Rodríguez L.; Herrera-Villarraga G.; Ochoa-Mogollón J.; Sarmiento-Sánchez C.; Rodríguez-Vargas L.H.; Rozo-Torres G.; Guayán-Ruíz P.; Sanjuan-Muñoz A.; Franco-Herrera A., Microplastics and plastisphere at surface waters in the Southwestern Caribbean sea, Journal of Environmental Manage |
| 2025 | Carrillo-Barragan et al. (2025) * | V4 16S rRNA; WGS | South Atlantic Ocean | 515f / 806r; WGS | Carrillo-Barragan P., Erni-Cassola G., Burkhardt-Holm P., Microbial Colonisation of Polyethylene in Offshore Marine Environments: Insights from the Southern and South Atlantic Oceans, (2025) |
